# PET imaging of fibroblast activation protein alpha (FAP) detects incipient cardiotoxicity due to anthracycline chemotherapy

**DOI:** 10.1101/2023.09.03.556130

**Authors:** Chul-Hee Lee, Onorina Manzo, Luisa Rubinelli, Sebastian E. Carrasco, Sungyun Cho, Thomas M. Jeitner, John W. Babich, Annarita Di Lorenzo, James M. Kelly

## Abstract

**Background:** Anthracycline chemotherapy is associated with a risk of cardiotoxicity leading to heart disease, particularly in pediatric cancer patients. Gold standard methods of detecting cardiotoxicity are insufficiently sensitive to early damage and specific pathophysiologies driving disease. Positron emission tomography (PET) couples anatomical resolution with biochemical mechanistic selectivity and potentially addresses the current diagnostic limitations in cardio-oncology. We aimed to validate PET imaging biomarkers targeting fibroblast activation protein alpha (FAP), Translocator protein (TSPO), and norepinephrine receptor (NET) for detection of incipient anthracycline-induced cardiotoxicity.

**Methods:** Cardiotoxicity was established in male C57BL/6J mice by a cumulative dose of 24 mg/kg doxorubicin (DOX) over 2 weeks. DOX mice and their age-matched controls were imaged with echocardiography and PET, using [^68^Ga]Ga-FAPI-04, [^18^F]DPA-714, and [^18^F]MFBG, over 12 weeks. Fractional shortening (FS) was determined from the echocardiograms, and cardiac uptake of the radioligands was quantified from the PET images. Heart sections were collected and used for the analysis of bulk RNA-seq, RT-qPCR, Western blot, in situ hybridization (ISH), and histopathological analysis.

**Results:** DOX mice exhibited cardiotoxicity and cardiac atrophy. Cardiac [^68^Ga]Ga-FAPI-04 PET signal was significantly higher in DOX mice from 2 weeks through the study endpoint. By contrast, no cardiac dysfunction was evident by echocardiography until 10 weeks, at which point FS was significantly reduced in DOX mice. There were no differences in [^18^F]DPA-714 and [^18^F]MFBG signals. Transcription and translation of FAP, but not TSPO or NET, was detected in cardiomyocytes and were elevated in the DOX hearts, in agreement with the PET data. Genes related to cell adhesion and extracellular remodeling were significantly upregulated in the DOX mice relative to controls.

**Conclusions:** FAP is a sensitive and selective imaging biomarker for incipient cardiotoxicity and FAPI PET is a promising non-invasive imaging tool for identifying patients at risk of cardiotoxicity during or after anthracycline chemotherapy.

**GRAPHICAL ABSTRACT:** 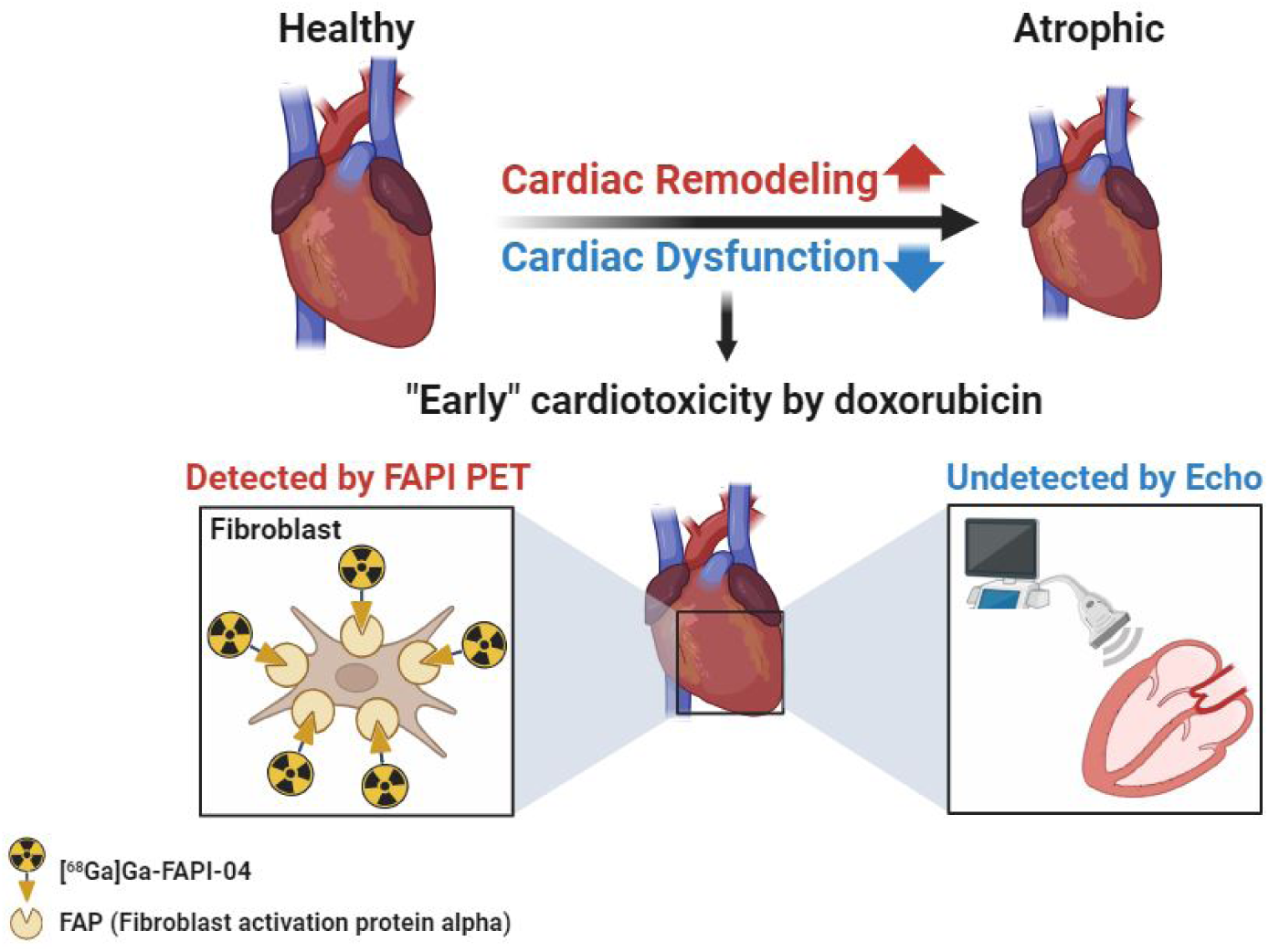

**NOVELTY AND SIGNIFICANCE:** *What is Known?:*

- Anthracycline chemotherapy results in cardiotoxicity for a sizeable population of treated patients. Cardiotoxocity manifests as cardiac dysfunction, and may result in long-term cardiac disease and heart failure, particularly in survivors of pediatric cancer.
- Cardiotoxicity is typically defined in terms of left ventricular ejection fraction (LVEF) deficits, as measured by echocardiography. However, this metric is often poorly sensitive to early disease and agnostic to underlying pathophysiology.
- Early treatment of cardiotoxicity improves recovery and long-term survival, emphasizing the need for accurate diagnostics in incipient disease.

*What New Information Does This Article Contain?:*

- [^68^Ga]Ga-FAPI-04 accumulates in the hearts of mice experiencing doxorubicin-induced cardiotoxicity as a function of fibroblast activation protein alpha (FAP) expression and activity. By contrast, cardiac uptake of radioligands targeting the translocator protein 18-kDa (TSPO) and the norepinephrine transporter (NET) do not differ between DOX animals and controls.
- Positron emission tomography (PET) imaging following administration of [^68^Ga]Ga-FAPI-04 detects abnormal cardiac remodeling significantly earlier than LVEF decrease is observed, indicating that it may be more sensitive to incipient disease. Our study identifies fibroblast activation protein alpha (FAP) as a promising diagnostic imaging biomarker in anthracycline-induced cardiotoxicity. We show that cardiac PET signal increases immediately after doxorubicin treatment, and the signal increase is sustained for at least 10 weeks. In addition, we demonstrate that FAP inhibitor (FAPI) PET correlates with expression of FAP protein and gene. Thus, we provide mechanistic insight into potentially-treatable pathophysiologies driving cardiac atrophy and toxicity, and have identified a translational PET tracer that can image the activation of these processes at an early stage.

## INTRODUCTION

Cancer therapy with the anthracycline doxorubicin (DOX) is the treatment of choice for a broad range of cancers, especially for the treatment of solid tumors and leukemias in adults and children.^1,2^ Roughly 60% of pediatric cancer patients today receive DOX as part of their treatment.^3^ However, despite being a mainstay of anti-cancer therapy, DOX can induce cardiovascular dysfunction, especially cardiotoxicity that results in heart failure.^4,5^ Heart conditions ranging from cardiomyopathy to heart failure are major adverse events in cancer patients treated with DOX, with up to 70% of total adverse events relating to cardiac health.^6,7^ Childhood cancer survivors are particularly susceptible, with more than 10% of those who received DOX treatment developing cardiotoxicity,^8^ which can develop into severe heart disease in adulthood.^9^ Early diagnosis of heart failure can lead to less invasive and more effective treatment. Therefore, the identification of emerging cardiotoxicity and attendant cardiac damage is an essential need to improve outcomes in cancer treatment.

Cumulative DOX exposure can produce the symptoms of cardiotoxicity within weeks or months (“early” cardiotoxicity) or after a number of years (“late”).^10^ The specific cellular and molecular changes responsible for these responses have not been fully elucidated.^11^ Assessment of left ventricular ejection fraction (LVEF) by echocardiography (echo) is currently the gold standard for evaluating cardiac function in patients with suspected cardiotoxicity.^12^ However, echo is subject to temporal variability^13–15^ and is poorly sensitive to early myocardial damage,^15,16^ while measurement of LVEF alone discounts other cardiopathologic effects that may occur.^17^ The introduction of tissue Doppler and strain imaging echo has allowed subclinical cardiotoxicity to be detected more reliably through speckle tracking-based deformation analysis.^10,18^ Nevertheless, the possibility of intervendor variability in strain measurements may require all follow up scans to be conducted using identical instrumentation.^18^ Furthermore, while improvements in echo image analysis have increased detection of subclinical cardiac functional decline, this method cannot be used to detect the underlying pathology responsible for these deficits. Cardiac magnetic resonance is an alternative imaging modality for assessing cardiac dysfunction,^19^ but this method still exhibits limited cost-effectiveness and uncertain diagnostic or prognostic value.^20,21^ Nuclear imaging methods, such as single photon computed tomography (SPECT), and positron emission tomography (PET), have been developed for assessing LVEF, myocardial viability, and perfusion^22^ but these do not provide mechanistic insight into cardiotoxicity. Circulating cardiac troponins and B-type natriuretic peptides are blood biomarkers of cardiac injury that can be assessed through minimally invasive procedures,^21,23^ however, their sensitivity and specificity may be insufficient to support use of these indices as single predictive biomarkers without further validation by an imaging technique,^5^ paticularly when samples are taken during early (acute) timepoints.^24^ To identify at-risk patients prior to anthracycline chemotherapy and detect incipient cardiotoxicity arising during or after chemotherapy therefore requires the validation of new biomarkers related to specific disease-causing pathophysiologies.

PET is a promising modality with which to pursue this aim. PET imaging can be combined with other modalities such as computed tomography (CT) to allow functional and anatomical imaging with spatial resolution as high as 2 mm.^25^ Recently, a number of radiolabeled small molecules for imaging fibroblast activation protein alpha (FAP),^26–29^ translocator protein 18-kDa (TSPO),^30–32^ and the norepinephrine transporter (NET)^33,34^ by PET have been described for detecting cancer-associated fibrosis, inflammation, and sympathetic innervation, respectively. Among these PET agents, FAP inhibitor [^68^Ga]Ga-FAPI-04,^27^ TSPO ligand [^18^F]DPA-714,^35^ and NET ligand *meta*-[^18^F]fluorobenzylguanidine, [^18^F]MFBG,^33^ have undergone preliminary clinical evaluation for oncologic and neurologic applications. Cardiac tissue remodeling,^36^ inflammation and cardiomyocyte mitochondrial dysfunction,^37^ and loss of cardiac sympathetic innervation^38^ are also identified as contributing pathologies in anthracycline-induced cardiotoxicity. Despite promising detection sensitivity, whole-body dosimetry, and relevance to key pathophysiologies of heart failure, neither these probes nor their molecular targets have been systematically evaluated in the context of cardiotoxicity.

Here, we established a preclinical model of DOX-induced cardiotoxicity for validating [^68^Ga]Ga-FAPI-04, [^18^F]DPA-714, and [^18^F]MFBG as diagnostic and prognostic biomarkers in cardio-oncology. Our goal was to evaluate the ability of these radioligands to detection incipient cardiotoxicity and identify the most suitable probe for translational to clinical imaging of cancer survivors treated with anthracyclines.

## METHODS

### General

Doxorubicin hydrochloride was purchased from Tocris Bioscience, USA and used without further purification. It was dissolved at a concentration of 0.75 mg/mL in sterile saline for injection (Hospira, USA) with the aid of sonication. The solution was stored in the dark at -20 °C for up to 24 h before use.

### Mouse model of doxorubicin-induced cardiotoxicity

All animal studies were approved by the Institutional Animal Care and Use Committee of Weill Cornell Medicine and were undertaken in accordance with the guidelines set forth by the U.S. Public Health Service Policy on Human Care and Use of Laboratory Animals. Adult male C57BL/6J (8-week-old) mice were purchased from The Jackson Laboratory, USA and randomly assigned to treatment (n=40) or control (n=16) groups. Mice in the treatment group were administered a solution of doxorubicin in saline at 3 mg/kg every other day for 2 weeks (total 8 doses; cumulative dose of 24 mg/kg) by intraperitoneal (i.p.) administration.^39^ Age-matched control (CTRL) mice were administered the same volume of saline i.p. The mice were weighed three times weekly and given access to food and water *ad libitum*.

### Echocardiography imaging and analysis

For evaluation of cardiac dimensions and function, echocardiography was performed under inhaled isoflurane anesthesia on a 37 °C heated platform using a Vevo 770 and 3100 Imaging systems (VisualSonics, Canada) accordingly to previously published methods.^40^ Briefly, scans were acquired using left-ventricle M-mode and all measurements were obtained by averaging the values of three consecutive cardiac cycles. Left-ventricle end-diastolic (LVDd) and end-systolic (LVDs) dimensions were measured using M-mode traces. Fractional shortening (FS) was calculated using the formula [(LVDd-LVDs)/LVDd]. Diastolic measurements were estimated at the point of maximum cavity dimension, and systolic were taken at the point of minimum cavity dimension, according to the American Society of Echocardiography’s recommended method.^41^

### Radiochemistry

[^68^Ga]FAPI-04,^27^ [^18^F]DPA-714,^30^ and [^18^F]MFBG^34^ were synthesized from their corresponding precursors according to literature procedures with minor modifications. Full experimental details describing the synthesis of the precursors and radioligands can be found in the Supplemental Material. All radioligands were formulated in 10% *v/v* ethanol/saline for administration to mice.

### Small animal microPET/CT imaging

DOX and CTRL mice were intravenously administered 100-150 µL of a 10% *v/v* ethanol/saline solution containing 3.7-11.1 MBq of the corresponding radioligand. The mice were imaged in groups of 2-4 using small-animal microPET/CT (Siemens Inveon™, USA) under isoflurane anesthesia (3.5% for induction, 1.5 % for maintenance) beginning 45 min after injection. The total PET acquisition time was 30 min, and a CT scan was obtained immediately before the PET acquisition for anatomic coregistration and attenuation correction. Images were reconstructed using the commercial Inveon software provided by the vendor. Images were corrected for decay and for the total activity injected.

### MicroPET/CT imaging data analysis

All microPET/CT images were evaluated with the AMIDE algorithm (A Medical Image Data Examiner).^42^ An ellipsoidal volume of interest (VOI) was generated for the heart and the right thigh muscle. The mean counts in the VOI were converted to percent injected dose per cubic centimeter (%ID/cm^3^) using the AMIDE algorithm, which was calibrated against a 1% injected dose standard. The %ID/cm^3^ in the heart was normalized against the %ID/cm^3^ in the muscle, providing a heart/muscle ratio, H/M.

### Preparation of heart tissue

The mice were anesthetized by i.p. ketamine injection and perfused with phosphate-buffered saline (PBS) via the left ventricle at a constant pressure of 80 mmHg. The hearts were patted dry and weighed on a digital balance. To perform the molecular and histological analysis, the hearts were cut transversally at the mid-horizontal plane. Cut fractions were separated for RNA and protein extraction, and tissue staining. Regions for tissue staining were fixed overnight in formalin at 4 °C and stored in 70% ethanol until further processing. The fractions for RNA and protein extraction were flash frozen in liquid nitrogen and stored at -78 °C until further use.

### Heart weight to tibia length (HW/TL) ratios

The tibia was collected ex vivo and all soft tissue was removed. The length was measured using a digital calipers. Heart weights were determined as described above and divided by the corresponding tibia length from the same animal to produce the heart weight to tibia length (HW/TL) ratio.

### Western blotting

Frozen heart tissue was pulverized by mortar and pestle for tissue homogenization. Ground heart tissue powders were collected and soaked in tissue protein extraction reagent (#78510, ThermoFisher, USA) supplemented with a protease inhibitor cocktail (#87786, ThermoFisher, USA) for protein extraction. Protein concentrations and western blot (WB) were performed as previously reported.^43^ The primary antibodies were anti-topoisomerase 2β) (TOP2β, 1:1000 dilution, #20549-1-AP, Proteintech, USA), anti-fibroblast activation protein alpha (FAP, 1:500 dilution, ab53066, Abcam, UK), anti-PBR (TSPO, 1:6000 dilution, ab109497, Abcam, UK), anti-SLC6A2 (NET, 1:250 dilution, MBS540046, MyBioSource, USA), and anti-HSP60 (1:1000 dilution, 12165S, Cell Signaling Technology, USA). The chemical luminescent signals were measured by Azure c400 Gel imaging system (Azure Biosystems, Inc. USA). Protein expression was quantified by drawing a region-of-interest (ROI) using ImageJ free software.

### Quantitative RT-PCR analysis

Ground heart tissue powders were collected and soaked in Trizol (Invitrogen, USA) and RNeasy Fibrous tissue mini kit (Qiagen, USA) was used to isolate total RNA from heart tissues. Genomic DNA was removed by DNase I (Qiagen), and RNA was reverse transcribed using an iScript kit (Bio-Rad, USA). The resulting cDNA was analyzed by quantitative RT-PCR (qPCR) using SYBR green master mix (Life Technologies, USA) on QuantStudio6 Real-Time PCR system (Life Technologies). mRNA levels were calculated by delta-delta CT method using the target gene (Fap) and reference genes (Rpl32, Tbp, Gapdh, and Actb). The full primer list is reported in **Table S5**.

### Bulk RNA-seq library construction and data analysis

The libraries were sequenced with paired-end 50 bps on the NovaSeq 6000 Sequencer (Illumina, USA). The raw sequencing reads in BCL format was processed through bcl2fastq 2.20 (Illumina) for FASTQ conversion and demultiplexing. After trimming the adaptors with cutadapt (version 1.18; https://cutadapt.readthedocs.io/en/v1.18/), RNA reads were aligned and mapped to the GRCm39 mouse reference genome by STAR (version 2.5.2; https://github.com/alexdobin/STAR),^44^ and transcriptome reconstruction was performed by Cufflinks (Version 2.1.1) (http://cole-trapnell-lab.github.io/cufflinks/). The abundance of transcripts was measured with Cufflinks using fragments per kilobase of transcript per million mapped reads (FPKM) as an output.^45,46^ Raw read counts per gene were extracted using HTSeq-count version 0.11.2.^47^ Gene expression profiles were constructed for differential expression, cluster, and principle component analyses with the DESeq2 package (https://bioconductor.org/packages/release/bioc/html/DESeq2.html).^48^ For differential expression analysis, pairwise comparisons were performed between two or more groups using parametric tests where read counts follow a negative binomial distribution with a gene-specific dispersion parameter. Corrected *p*-values were calculated based on the Benjamini-Hochberg method to adjust for multiple testing.

For the differentially expressed genes (DEGs) analysis, *p* < 0.01 was used as the signifier of statistical significance, and Log2FC (FC, fold change) ≥ 0.55 and Log2FC ≤ -0.85 were used to distinguish upregulated (Up) and downregulated (Down) DEGs, respectively. The heat map was generated using R studio to compare DEGs between groups, and the volcano plot for the overall distribution of DEGs was analyzed using GraphPad Prism 9.0 (GraphPad Software, USA).

### DAVID analysis and establishment of PPI networks

The database for annotation, visualization, and Integrated Discovery (DAVID) was used to group DEGs based on biological function (https://david.ncifcrf.gov/). The 1326 Upregulated genes and 1684 Downregulated genes were submitted for the Gene Ontology (GO) according to the biological process (BP) analysis. The heat map for the GO:BP data was generated using R studio. A protein-protein interaction (PPI) network was developed to identify the association between a target and related DEGs by utilizing the STRING database (http://string-db.org/).^49^ GO terms and PPI networks with a *p-*value cutoff < 0.05 were regarded as significant.

### Histopathology

The tissue was processed in alcohol and xylene and embedded in paraffin. Four transverse sections of the heart per mouse, including right and left ventricles, right and left auricles, and interventricular septum were sectioned at 5-µm thickness and stained with hematoxylin and eosin. Histopathological evaluation of the heart was performed by a board-certified veterinary pathologist. Hearts were evaluated on the basis of cardiomyocytes showing necrosis, degeneration (cytoplasmic vacuolization), and atrophy, leukocytic cell infiltrates, and interstitial fibrosis.

Formalin fixed sections of the heart were stained with Masson’s Trichrome to evaluate the presence of collagen in cardiac tissues. To determine the percentage of collagen in the heart, digital whole slide images of Masson’s trichrome-stained hearts were manually annotated and then classified pixels were evaluated with a random forest algorithm using QuPath (an open-source software for digital pathology image analysis accessed through: https://qupath.github.io/). Regions of collagen for this analysis included collagen fibrils between cardiomyocytes and around preexisting vasculature within cardiac musculature. Regions excluded for this analysis included preexisting collagen from great vessels, leaflet insertion bands, and pericardial connective tissue.

### Immunohistochemistry

Formalin-fixed, paraffin-embedded sections were stained using an automated staining platform (Leica Bond RX, Leica Biosystems). Following deparaffinization and heat-induced epitope retrieval in a citrate buffer at pH 6.0, the primary antibody against TSPO, also known as peripheral-type benzodiazepine receptor (PBR; ab109497, Abcam, Waltham, MA), was applied at a dilution of 1:10000. A rabbit anti-goat secondary antibody (Cat. No. BA-5000, Vector Laboratories, Burlingame, CA) and a polymer detection system (DS9800, Novocastra Bond Polymer Refine Detection, Leica Biosystems) was then applied to the tissues. The chromogen used was 3,3’-diaminobenzidine tetrachloride (DAB) and the sections were counterstained with hematoxylin and examined by light microscopy. Positive immunoreactivity for TSPO was confirmed with internal mouse tissue array controls used to validate this immunoassay. A subset of tissues incubated with antibody diluents and secondary antibody only were used as negative controls for this assay. Images were acquired with an Olympus VS200 slide scanner (Olympus, Tokyo, Japan) with a 20x objective. Quantitative image analysis was performed by using the QuPath Pixel classifier module. A random forest algorithm was used for identifying pixels as TSPO-positive, TSPO-negative and background in cardiac sections. Region of interest and thresholding values were validated by a board-certified veterinary pathologist.

For CD11b immunohistochemistry, a heat-mediated antigen retrieval with citrate buffer (pH 6.0) was applied on deparaffinized cardiac sections, which were then incubated with a primary anti-CD11b antibody at a dilution of 1:4000 (ab133357, Abcam, USA). A goat anti-rabbit secondary antibody (Cat. No. BA-1000, Vector Laboratories) and a polymer detection system (DS9800, Novocastra Bond Polymer Refine Detection, Leica Biosystems) were then applied to the tissues. The chromogen was DAB, and the sections were counterstained with hematoxylin and examined by light microscopy.

### In Situ Hybridization

Formalin-fixed, paraffin-embedded cardiac sections were incubated with the target probe designed to detect region 486 - 1588 of murine fibroblast activation protein (Fap) mRNA, NCBI Reference Sequence NM_007986.3 (RNAscope® LS 2.5 probe for murine FAP, #423888; Advanced Cell Diagnostics, Newark, CA). The target probe was validated on sections of murine skin and heart from mice. Slides were stained on an automated stainer (Leica Bond RX, Leica Biosystems) with RNAscope 2.5 LS Assay Reagent Kit-Red (322150, Advanced Cell Diagnostics) and Bond Polymer Refine Red Detection (DS9390, Leica Biosystems). Control probes detecting a validated positive housekeeping gene (mouse *peptidylprolyl isomerase B*, *Ppib* to confirm adequate RNA preservation and detection; 313918, Advanced Cell Diagnostics) and negative control, *Bacillus subtilis* dihydrodipicolinate reductase gene (*dapB* to confirm absence of nonspecific labeling; 312038, Advanced Cell Diagnostics) were used. Positive RNA hybridization was identified as discrete, punctate chromogenic red dots under bright field microscopy. Images were acquired with an Olympus VS200 slide scanner with a 40x objective. Quantitative image analysis of Fap hybridization was performed with QuPath using an algorithm for singleplex chromogenic RNAscope image analysis. Fap positive hybridization signal was classified as follows: 1 red dot / cell, 2 red dot / cell, and 3+ red dots / cell in each transverse section of the heart. An H-score of Fap positive signal from each sample was calculated by the QuPath software. Samples with autolysis or regions in the tissue with pale-brown precipitate and/or folding artifacts were excluded from this analysis.

### Statistical analysis

Statistical analyses were carried out using GraphPad Prism 9.0. All data were expressed as means ± standard deviation (SD) and are representative of at least three separate biological experiments. The unpaired two-tailed Student’s *t-test* or Mann-Whitney test was determined for comparisons of two groups. For correlation analysis, the Pearson correlation test was used. A *p-*value of less than 0.05 was considered statistically significant.

## RESULTS

### DOX treatment in mice induces cardiotoxic physiological and molecular changes

To establish a clinically-relevant model of DOX-induced cardiotoxicity, we administered intraperitoneal saline (control; CTRL) or DOX (cumulative dose 24 mg/kg) over 2 weeks (W)^39^ to C57BL/6 mice followed by 10-12 W observation together with serial echo and microPET/CT imaging **(Figure 1A)**. In agreement with multiple literature reports,^11,24^ we observed significantly lower body weights in the DOX mice compared to the age-matched CTRL group **(Figure 1B; Table S1)**. The heart weight (HW) to tibia length (TL) ratio was 40% lower in DOX groups compared to the CTRL group at from 7 to 12 W *(p* < 0.0001) **(Figure 1C; Table S2)**. There were not significant differences in TL between the two groups *(p* = 0.5566) **(Figure S1A)**, indicating that cardiac atrophy was occurring in the DOX animals, as previously reported.^11^ Next, to evaluate the effect of DOX treatment at a cellular level, the expression of TOP2β, a primary mediator of DOX-induced toxicity,^50^ was evaluated. We observed rapid and sustained decrease in TOP2β expression, which persisted up to 10 W **(Figure 1D)**. Collectively, these experiments confirmed that a cumulative dose of 24 mg/kg DOX induced sustained cardiac atrophy and reduction of TOP2β protein expression in mice.

**Fig. 1.**
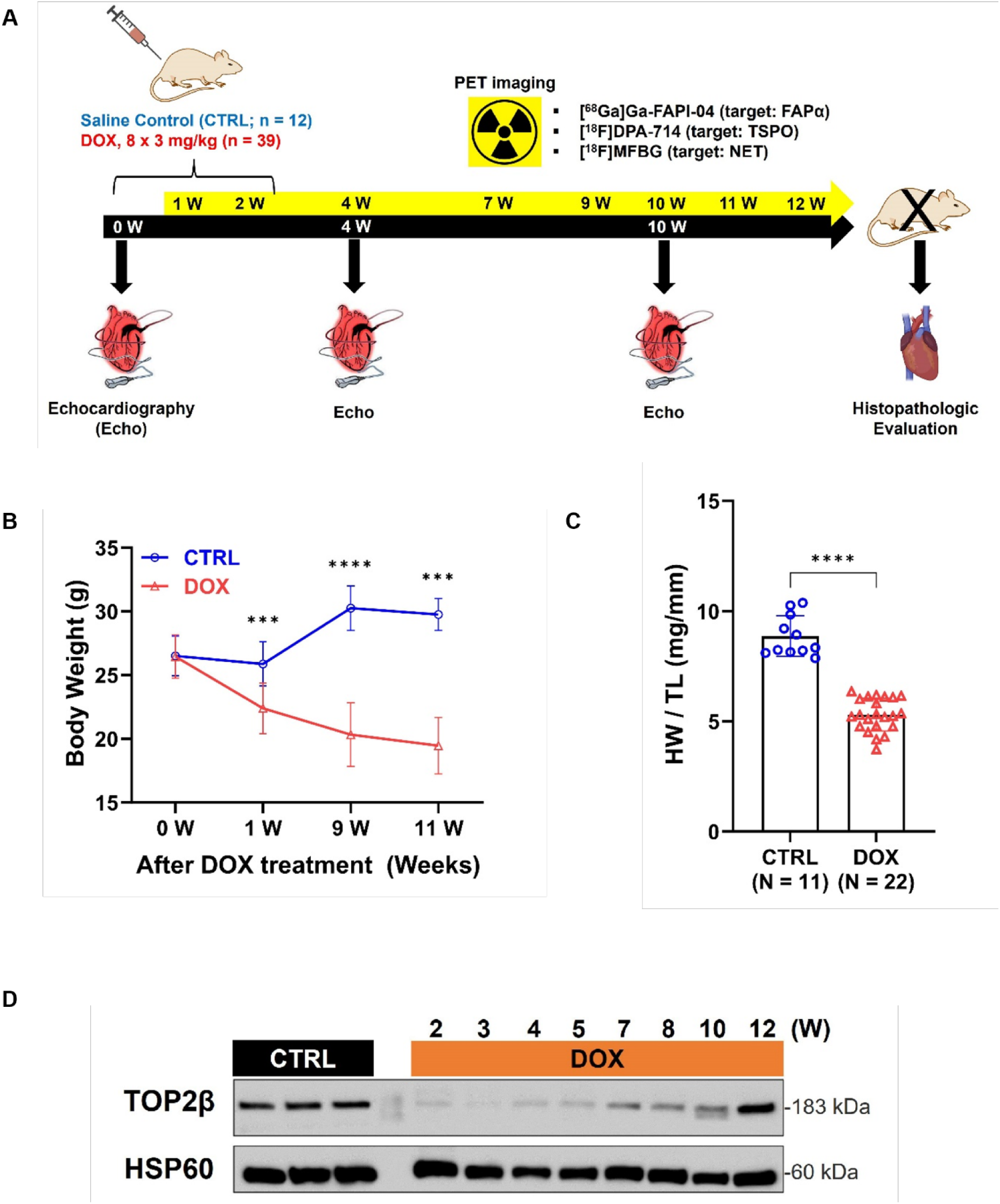
Establishment of clinically-relevant cardiotoxic mouse model induced by systemic administration of DOX. **(A)** Schematic of this study. After establishing the DOX-induced cardiotoxicity model, underlying pathophysiology was evaluated by serial PET imaging using [^68^Ga]Ga-FAPI-04 (for targeting FAP), [^18^F]DPA-714 (for targeting TSPO), and [^18^F]MFBG (for targeting NET). Echocardiography (echo) imaging was performed at 4 weeks and 10 weeks. Heart tissue was extracted at the PET imaging time points for measurement of heart weight and evaluation of molecular and cellular changes in cardiac tissue. **(B)** Body weights were compared between CTRL and DOX groups over the course of the experiment. **(C)** Heart weight indexed to tibia length. **(D)** TOP2β expression from the heart lysates was evaluated by western blot. HSP60 was used as an internal control. Data are presented as the mean ± s.d. *** *p < 0.001* and **** *p < 0.0001*.

### [^68^Ga]Ga-FAPI-04 PET detects cardiac abnormalities earlier than echo

We performed echo imaging and analysis in CTRL and DOX groups at acute (4 W) and chronic (10 W) phases.^24^ At 4 W, left-ventricle end-diastolic diameter (LVDd) and left-ventricle end-systolic diameter (LVDs) were not significantly different between the CTRL (n = 6) and DOX (n = 6) animals (LVDd; *p* = 0.833, LVDs; *p* = 0.165) **(Figure 2A-C)**. However, LVDs was significantly increased by 20% in the DOX group (n = 5) at 10 W *(p* < 0.01) **(Figure 2A and C)**. In parallel, we observed no significant change in LVDd *(p* = 0.149) **(Figure 2B)**. As a result, fractional shortening (FS) decreased from 40% in the CTRL animals and DOX animals at 4 W to less than 30% at 10 W *(p* < 0.01) **(Figure 2D)**. At this later time point, blood cardiac troponin-I (CTNI) levels were higher in DOX mice, but the difference was not significant *(p* = 0.24*7)* **(Figure S2)**.

**Fig. 2.**
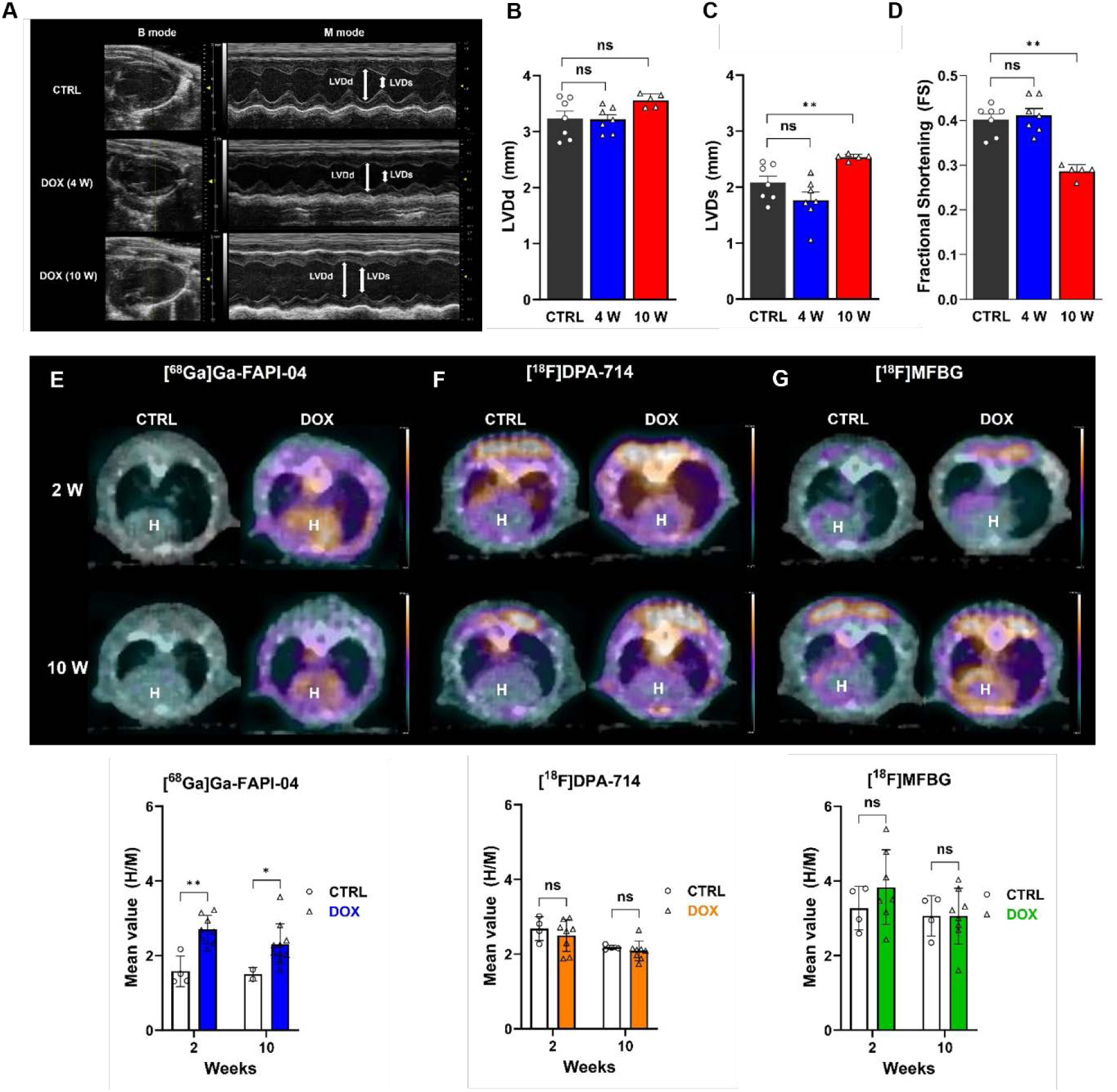
[^68^Ga]Ga-FAPI-04 PET detects cardiac remodeling before it is evident by conventional echo. Pathology was monitored by echo and microPET/CT imaging. DOX-treated mice were randomly assigned to imaging groups for comparison with age-matched controls (CTRL). **(A)** Representative images of the left ventricle (LV) serial echo of CTRL (n =7) and DOX groups (4 W, n = 7 and 10 W, n = 5). Left ventricular end-diastolic diameter (LVDd) **(B)** and left ventricular end-systolic diameter (LVDs) **(C)** were determined from the echo scans. **(D)** Fractional shortening (FS) was calculated from LVDd and LVDs. Representative [^68^Ga]Ga-FAPI-04 **(E)**, [^18^F]DPA-714 **(F)**, and [^18^F]MFBG **(G)** microPET/CT fusion transaxial images at 2 and 10 W. Quantitative [^68^Ga]Ga-FAPI-04, [^18^F]DPA-714, and [^18^F]MFBG PET signals (mean value of % injected dose per centimeter cubed, %ID/cm^3^) was normalized by thigh muscle uptake. The number of mice imaged at each time point is displayed in **Table S3**. Data are presented as the mean ± s.d. * *p < 0.05,* and ** *p < 0.01*.

We performed serial microPET/CT imaging with [^68^Ga]Ga-FAPI-04, [^18^F]DPA-714, and [^18^F]MFBG for 12 W to compare the time course of cardiac uptake differences with the time course of functional deficits in the DOX hearts **(Figure S3; Table S3)**. To account for differences in heart function, we normalized cardiac uptake to skeletal muscle (thigh muscle). Skeletal muscle has minimal basal expression of our molecular targets and therefore acts as a surrogate for blood pool effects. Furthermore, it is also subject to DOX-induced toxicity^11^ and therefore controls for off-target effects. [^68^Ga]Ga-FAPI-04 uptake was significantly increased in the acute phase (2 W; 1.7-fold) *(p* < 0.01) when no evidence of cardiotoxicity is evident as measured by echo **(Figure 2E).** Moreover, increased cardiac [^68^Ga]Ga-FAPI-04 uptake persisted in the DOX mice through the chronic phase (10 W; 1.5-fold) *(p* < 0.05) (**Figure 2E, Figure S3**). By contrast, there was no significant difference between cardiac uptake of [^18^F]DPA-714 and [^18^F]MFBG in the CTRL and DOX groups at either the early or late phases **(Figure 2F and G, Figure S3).**

### Increased cardiac [^68^Ga]Ga-FAPI-04 uptake is significantly correlated to FAP expression at the gene and protein levels

Next, we sought to validate the significant differences in cardiac [^68^Ga]Ga-FAPI-04 uptake through determining their correlation with FAP protein and mRNA expression. Cardiac tissue was collected from perfused hearts at 4 W for Western blot, qPCR, and RNA-seq analyses and tissue staining **(Figure 3A)**. As expected, FAP expression was 2.9-fold higher *(p* < 0.01) in DOX mice than CTRL animals, while TSPO and NET showed no significant differences in protein expression **(Figure 3B and C)**. At the same time, we determined the Fap gene expression using three different Fap primers and different reference genes to ensure accurate qPCR analysis in spite of the heterogeneity of our heart tissue **(Table S4)**.^51^ Fap gene expression increased 2.5-fold (normalized to Rpl32, *p* < 0.0001), 2.0-fold (normalized to Tbp, *p* < 0.0001), and 4.7-fold (normalized to Gapdh, *p* < 0.0001), respectively, compared with CTRL animals **(Figure 3D)**. In agreement with our [^68^Ga]Ga-FAPI-04 PET imaging, cardiac Fap gene expression in DOX mice increased 1.7-fold (*p* < 0.01), 2.8-fold (*p* < 0.0001), and 2.8-fold (*p* < 0.0001) relative to CTRL mice at 2, 7, and 10 W, respectively **(Figure S4A)**. We observed a similar trend in FAP protein expression (**Figure S4B**), suggesting that mRNA and protein expression levels are proportional in this model. Additionally, FAP activity was significantly increased in DOX hearts compared to controls (*p* = 0.031) (**Figure S5**).

**Fig. 3.**
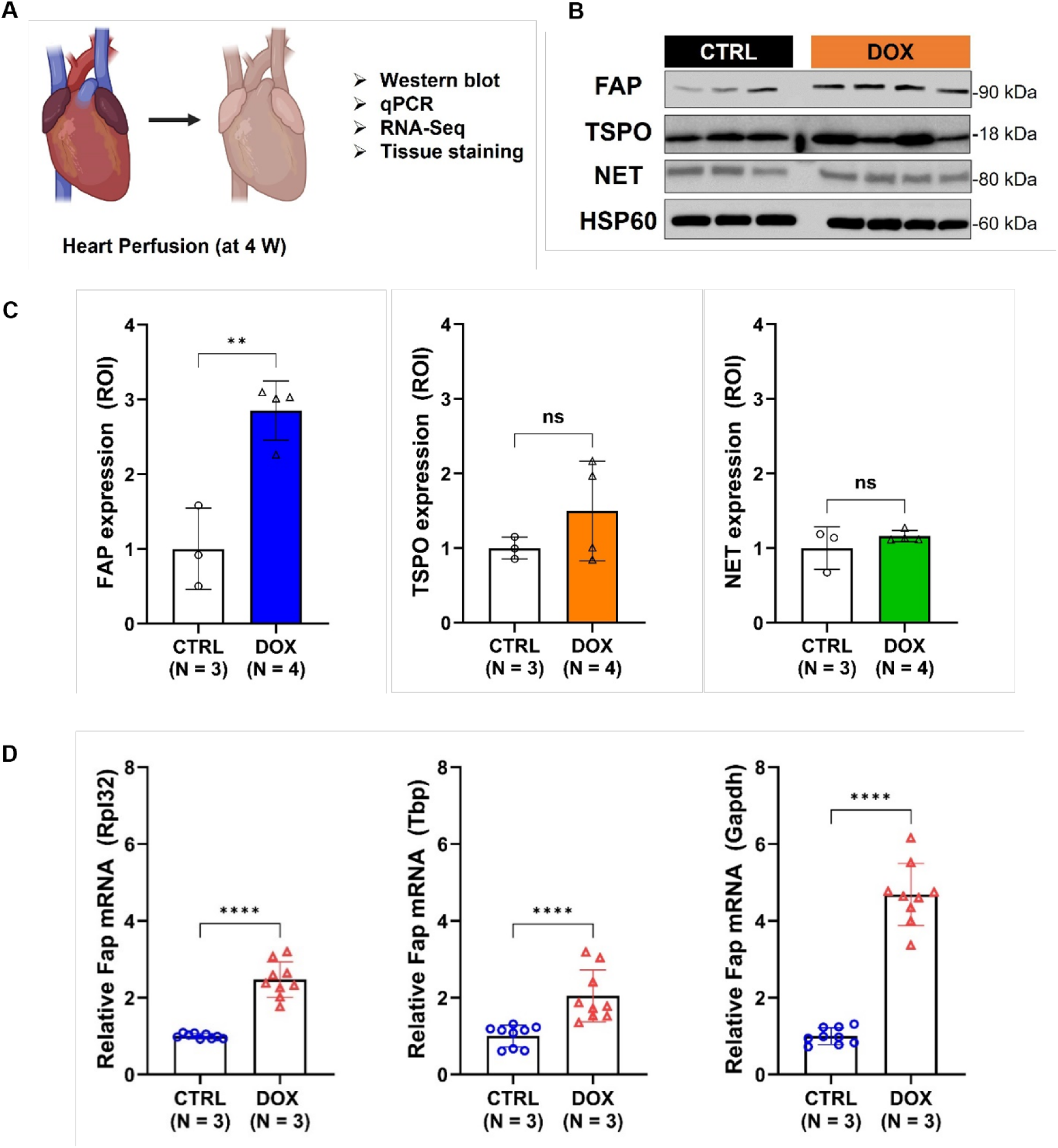
Expression of FAP protein and mRNA is elevated at 4 W. **(A)** Isolation of heart tissues for determination of mRNA, protein, and tissue levels. **(B)** Protein expression of FAP, TSPO, and NET (CTRL; n = 3, DOX; n = 4) was evaluated by western blot. HSP60 was used as an internal control. **(C)** Each band size was normalized by drawing ROIs using ImageJ free software. **(D)** RT-qPCR (CTRL; n = 3, DOX; n = 3) was performed for the validation of Fap mRNA expression using three different primers, listed in **Table S5**. Rpl32, Tbp, and Gapdh were used as reference genes in cardiac tissue. Data are presented as the mean ± s.d. ** *p < 0.01* and **** *p < 0.0001*. The schematic illustration (A) was drawn using https://biorender.com/

### FAP is a diagnostic imaging biomarker for detecting incipient cardiotoxicity by PET

Having established that the cardiac PET signals of our candidate probes correlated with protein and mRNA expression of the corresponding molecular target, we sought to validate uptake by tissue staining. We separated a cohort of DOX animals (n = 9) into high or moderate uptake groups **(Figure 4A)**. Cardiac [^68^Ga]Ga-FAPI-04 PET signal was significantly higher in these animals than in the age-matched CTRL mice (n = 4; *p* < 0.05). Interestingly, DOX-treated mice showed little evidence of pathological changes in cardiomyocytes. A mild degree of individual cardiomyocyte necrosis and degeneration and/or focal to multifocal aeas of myocardial fibrosis were occasionally observed in four DOX hearts. The rest of the DOX hearts (n = 11) did not show pathological changes.

**Fig. 4.**
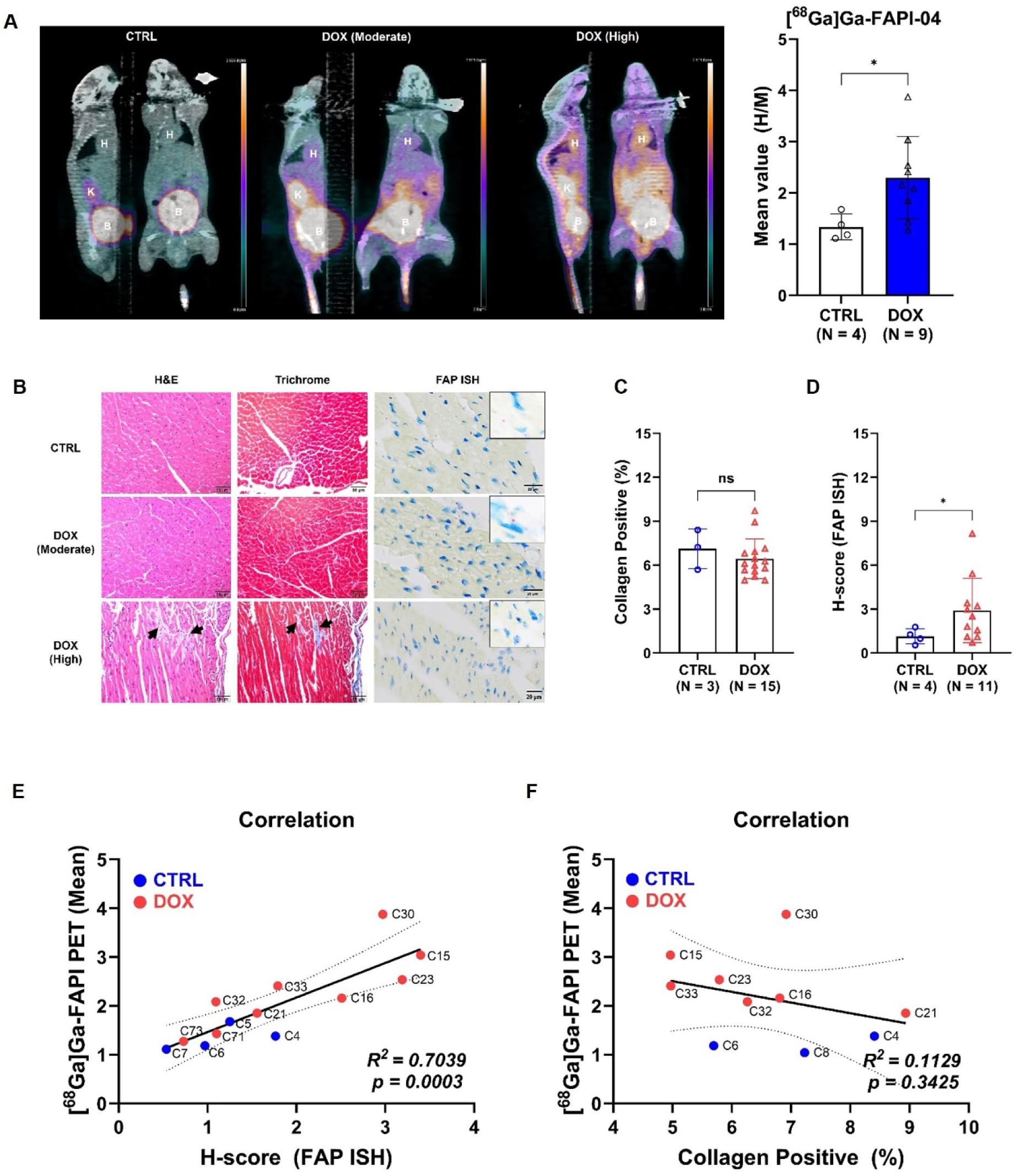
Correlation between FAPI PET, FAP in situ hybridization (ISH), and trichrome staining. **(A)** *Left*: Representative whole-body FAPI PET sagittal and coronal maximum intensity projections (MIPs) acquired 45 min post injection of [^68^Ga]Ga-FAPI-04. DOX mice showed moderate or high uptake compared to CTRL. *Right*: Image-based quantitation of cardiac PET signal normalized by thigh muscle uptake (H/M). **(B)** Representative H&E, Trichrome, and FAP ISH stains. Myocardial fibrosis was occasionally and regionally detected in DOX-treated hearts (arrows). Fap mRNA (punctate red dots) was detected in the cytoplasm and/or nuclei of cardiomyocytes and/or epicardial stromal cells (Insets). **(C)** Comparison of collagen positive regions from the trichrome staining. Quantitation was performed using QuPath software, with positive regions defined as deposition of collagen fibril between cardiomyocytes. **(D)** Comparison of the H-score from FAP ISH staining. Quantitation was performed using QuPath software. **(E)** A significant correlation was observed between [^68^Ga]Ga-FAPI-04 cardiac PET signals and FAP expression in the corresponding tissues *(p = 0.0003)*. **(F)** No correlation was observed between [^68^Ga]Ga-FAPI-04 cardiac uptake and the percentage of collagen positive regions in the corresponding tissues *(p = 0.3425)*. Data are presented as the mean ± s.d. * *p < 0.05.* Scale bar: 1 mm and 20 μm.

As DOX associated heart damage can lead to fibroblast activation and subsequent interstitial fibrosis, the percentage of collagen was evaluated by the use of Masson’s Trichrome staining of the hearts. There were no differences in H&E and Masson’s trichrome staining between these three groups **(Figure 4B)**. Indeed, collagen-positive regions averaged approximately 7% for CTRL mouse hearts (n = 3) and approximately 6.5% for DOX hearts (n = 15) **(Figure 4C)**. However, spatiotemporal Fap expression, as determined by in-situ hybridization (ISH), was significantly higher in the DOX animals (n = 11; *p* < 0.05) **(Figure 4D)**. Fap nucleic acid was detected in the cytoplasm and associated with the nuclei of cardiomyocytes and stromal cells. Furthermore, the H-score was higher in the tissue slices belonging to mice in the high uptake group than mice in the moderate uptake group **(Figure 4B; Table S6)**. There was a linear correlation between [^68^Ga]Ga-FAPI-04 PET signal and H-score (*p* < 0.001) **(Figure 4E)** but no correlation between PET signal and collagen formation *(p* = 0.343) **(Figure 4F)**. As predicted by our PET imaging, there was no difference in TSPO staining between DOX and CTRL mice **(Figure S6)**. We observed no NET staining in either CTRL or DOX samples (*data not shown*). Taken together, these results indicate that FAPI PET is a potential diagnostic biomarker in the DOX model.

### DOX promotes cardiac remodeling and disrupts mitochondrial energetics

To investigate the role that FAP may be playing in DOX-induced cardiotoxicity, we performed bulk RNA-seq analysis. We first constructed a volcano plot using Log2FC and a negative Log False discovery rate (FDR) with 14,101 DEGs. Although none of the Fap, Tspo, and Slc6a2 genes showed significant differences in the overall DEGs population, Fap gene expression did significantly increase when the expression level was normalized by fragments per kilobase of transcript per million (FPKM) mapped fragments **(Figure S7A and B)**. We also generated a heat map of all DEGs. Clear differences between the DOX and CTRL groups were evident, though we observed a degree of heterogeneity within each group **(Figure 5A)**. Compared with CTRL mice, DOX hearts showed 1326 markedly upregulated genes and 1684 markedly downregulated genes. These genes were used to identify the top 20 from the *p-*value affected biological processes (BP) in the gene ontology (GO). Within the upregulated genes, the most significantly affected GO:BPs included those related to intracellular signal transduction, protein phosphorylation, cell adhesion, angiogenesis, and extracellular matrix organization. Within the downregulated genes, on the other hand, the most affected GO:BPs were those related to mitochondrial translation, ATP synthesis, and respiratory chain complex **(Figure 5B)**. Furthermore, numerous GO:BPs associated with cardiac remodeling were identified in the upregulated group, whereas GO:BPs associated with mitochondrial energetic dysfunction were identified in the downregulated group **(Figure S8)**. Taken together, these data indicate that DOX treatment induces cardiac remodeling and impairs mitochondrial energetics in cardiomyoctyes.

**Fig. 5.**
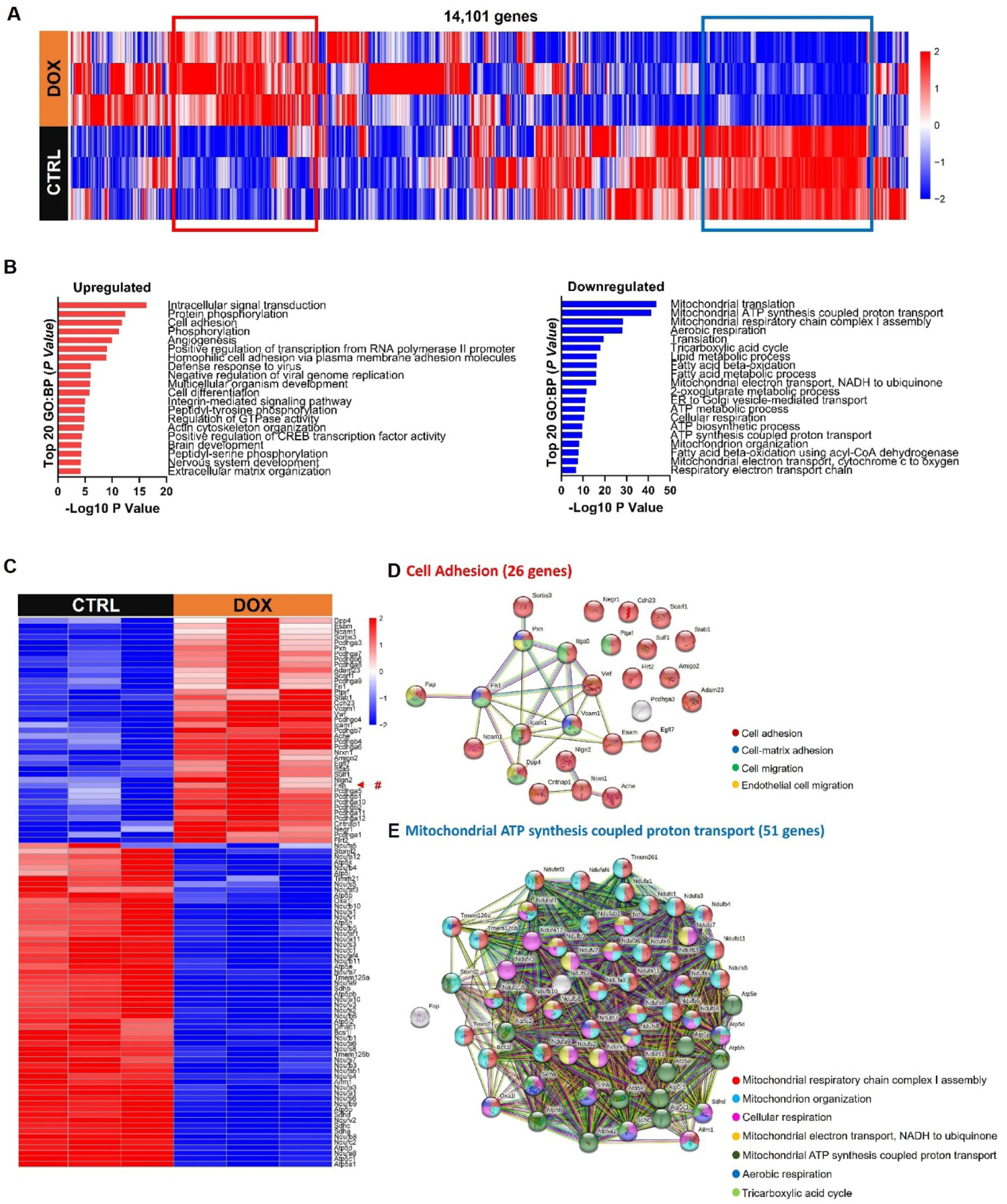
The apparent difference of biological processes in gene ontology according to DOX treatment. **(A)** Heat map representation of the 14,101 genes from the same samples of qPCR. **(B)** The top 20 up-(red) or downregulated (blue) biological processes (BP) of gene ontology (GO) by *P* value were determined from the bulk RNA-seq data. **(C)** Heat map representation from the GO:Cell adhesion (upregulated; 26 genes) and GO:Mitochondrial ATP synthesis coupled proton synthesis (downregulated; 51 genes) with Fap (marked by #). **(D and E)** STRING database highlighting interactions between Fap and highly upregulated or downregulated genes.

The heat map was refined by selecting representative genes from upregulated or downregulated one BP of the top 3 GO:BP terms, including the Fap gene, and excluding genes with heterogeneous expression **(Figure 5C)**. This heat map highlights the increased expression of Fap (indicated by the hashtag) in DOX hearts compared to CTRLs. Next, we analyzed the same gene family using the STRING database. FAP strongly interacted with fibronectin-1 (Fn1), a major component of cardiac ECM remodeling and fibrosis,^52,53^ which in turn associated with other proteins involved in cell adhesion and migration **(Figure 5D)**. On the other hand, there was no protein association between FAP and any component of the downregulated group **(Figure 5E)**. From these findings, we conclude that increased FAP expression contributes to multiple processes involved in cardiac remodeling but does not predict fibrosis in cardiac tissue.

## DISCUSSION

In this study, we sought to validate PET imaging targets that are more sensitive to the early symptoms of DOX-induced cardiotoxicity than conventional echo and are biomarkers of specific pathophysiologies. Nuclear medicine approaches have been underutilized for this purpose due to concerns about radiation exposure and lack of widespread availability of imaging devices in cancer treatment facilities. Nevertheless, this imaging technique allows anatomical information to be coupled to biochemical information afforded by a suitably chosen probe targeting a specific disease-relevant molecule or pathway. Although DOX-induced cardiotoxicity ultimately decreases LVEF, compromises cardiac performance, and can lead to heart failure, the mechanisms by which it does so may vary between patients. Non-invasive detection of these mechanism by molecular imaging is a safe and important way of assessing individual-to-individual variations. This is significant because the appropriate cardioprotective treatment could be administered on an individual basis with suitable knowledge of the specific underlying pathology. This is the premise behind personalized medicine.

The primary nuclear medicine approaches to imaging cardiotoxicity focus on imaging perfusion.^54^ Recently, new probes have been developed to image specific pathophysiologies, including mitochondrial damage,^55,56^ sympathetic inervation,^34,57–59^ inflammation,^31,60–62^ cardiac remodeling,^63–67^ and cardiac metabolic dysfunction.^68–70^ These probes have largely been studied in preclinical models, where they have provided important insight into the development and progression of cardiotoxicity. To build on this work, we identified cardiac remodeling, inflammation, and inervation as plausible contributors to DOX-induced cardiotoxicity that could arise at the earliest stages of disease. We selected three radioligands, [^68^Ga]Ga-FAPI-04,^71,72^ [^18^F]DPA-714,^35,73^ and [^18^F]MFBG,^74^ that are already under clinical investigation for other indications to image these processes. Preliminary reports indicate that the radiation dosimetry of these probes is acceptable, thereby supporting their use in cancer survivors. We anticipate that this will facilitate the future clinical translation of our PET approach.

To further accelerate clinical translation, we applied a well-established model of cardiotoxicity encompassing subacute and chronic phenotypes.^24,39^ This heterogenous model reflects the reality that anthracycline-induced cardiotoxicity is a complex process that involves multiple cell types in heart tissue^75^ and often differs between even those patients receiving the same dose of anthracycline. In cardiomyocytes with high TOP2β expression,^76^ DOX binds to DNA form a complex with TOP2β that triggers cell death pathways.^5,50^ Consequently, cancer patients with high levels of TOP2β in cardiomyocytes are likely to be more susceptible to DOX-induced cardiotoxicity.^50,77^ Our Western blot data confirm that TOP2β expression is rapidly downregulated in response to DOX treatment, with gradual recovery over the 12 W observation period. Furthermore, we observed body weight decrease and diminishing HW/TL ratios after DOX treatment, consistent with established models of DOX-induced cardiotoxicity^11,78^. Although not statistically significant at this dose, heart weight was a lower percentage of body weight in the DOX mice than the controls (**Figure S1B**), suggesting that cardiac atrophy is more pronounced than cachexia or other systemic effects in these animals.^11^ Finally, functional declines in cardiac performance emerged by 10 W, even when corresponding interstitial fibrosis was not widely observed. These observations indicate a spectrum of symptoms consistent with clinical presentation of cardiotoxicity.

Our transcriptomic data indicate that cardiac remodeling is initiated in response to DOX treatment. Pathological cardiac remodeling arises in chronic heart failure through the activation of multiple pathways,^79,80^ and we observed intracellular signaling,^81–83^ cell adhesion,^84^ angiogenesis,^85^ extracellular matrix remodeling,^86,87^ and cell migration,^88^ to be highly enriched in our DOX tissues. Another major aspect of remodeling is cardiac fibrosis. This process is typically initiated by activated cardiac fibroblasts and eventually leads to the functional change of heart tissue and diastolic dysfunction.^89^ FAP is a marker of activated cardiac fibroblasts^89^ and has recently been targeted by radiopharmaceuticals for PET imaging in cardiovascular disease and heart failure.^65,90–92^ In addition, a recent case study speculated that incidental cardiac FAPI PET signal detected in a cancer patient may have been due to cardiotoxicity arising from the chemotherapy regimen,^7^ although this hypothesis was not explored further. Furthermore, the BioGPS^93^ and GTEx databases indicate basal Fap gene expression to be moderate in cardiac tissue of mice and humans (**Figure S9)**, which may enable relatively small changes in expression to be detected by molecular imaging. These observations provided the rationale for our hypothesis that FAPI PET would detect incipient cardiotoxicity.

We demonstrate that the cardiac signal intensities of [^68^Ga]Ga-FAPI-04 PET increased almost immediately after DOX treatment, substantially earlier than any functional alteration could be imaged by echo. This increased signal, which was sustained throughout the 12 W observation period, significantly correlated with expression of FAP protein and Fap mRNA. We were unable to successfully perform immunohistochemistry for murine FAP using commercially available antibodies, and therefore used ISH to assess the distribution of FAP in cardiac tissue. Our experiments confirm the specificity of FAPI PET for DOX-induced alterations in cardiac FAP, thereby highlighting the initiation of pathological remodeling pathways in the injured heart. We also provide a mechanistic link between elevated PET signal and disease, an outcome that has not been accomplished in human patients due to inconclusive histopathology studies.^65,66^ Significantly, our data indicate that FAP is a diagnostic imaging biomarker in cardiotoxicity that might be superior to echo in detecting early mediators of cardiac damage.

Our study did not conclusively establish whether FAP is also a prognostic imaging biomarker in cardiotoxicity. We did not find a correlation between PET signal and fibrosis in our tissue samples. However, fibrosis was minimal in our model. We conclude that FAP in this model participates more broadly in cardiac remodeling. Consistent with literature reports in mice and humans,^11,94^ DOX treatment induced cardiac atrophy in our mice. Cardiac atrophy requires extensive remodeling of the ECM due to loss of cardiomyocyte mass,^95^ consistent with the upregulation of ECM remodeling pathways evident in our RNA-seq analysis (**Figure 5B; Figure S7**). We did not see evidence of substantial cardiomyocyte death, which may explain why we observed minimal fibrosis in this model. Indeed, reports of fibrosis in atrophied hearts are conflicting,^94^ which likely reflects both the techniques used to quantify collagen and the prevalence of remodeling pathways that do not result in collagen deposition. Nevertheless, given that DOX induces cardiotoxicity through a variety of molecular mechanisms,^96^ it is possible that FAP PET might correlate with other indices of disease severity. For example, we observed a negative correlation between cardiac FAPI PET signal and HW/TL at the end of the study (**Figure S10**). We will need larger group sizes, longer follow up periods, alternative indices of disease severity, and perhaps more acute pathology to determine if elevated cardiac FAPI PET signal corresponds to more severe long-term outcomes.

Our alternative molecular targets, TSPO and NET, proved to be neither diagnostic nor prognostic biomarkers in DOX-induced cardiotoxicity. Our rationale for targeting TSPO was the prominent role that oxidative stress and inflammation play in DOX-induced cardiotoxicity.^37^ TSPO is expressed not only in cardiomyocytes^97^ but also in activated immune cells, especially macrophages.^98–100^ Resident and circulating macrophages are implicated in the response to DOX-induced cardiotoxicity.^101^ Preclinical studies have demonstrated increased cardiac uptake of [^18^F]DPA-714 in mice with inflammatory heart conditions,^60,102^ but we observed neither a significant increase in [^18^F]DPA-714 signal nor an increase in TSPO staining by immunohistochemistry. Moreover, we found no evidence of macrophage infiltration by histology or CD11b immunohistochemistry **(Figure S11**). TSPO is also present in the mitochondrial outer membrane of cardiomyocytes where it modulates oxidative stress and regulates mitochondrial physiology and metabolism.^103^ We observed downregulation of a number of genes related to mitochondrial metabolism in the DOX mice, but this did not translate to increased [^18^F]DPA-714 uptake. Given the nearly ubiquitous expression of TSPO in tissue,^104^ it is possible that substantial off-target uptake reduces the sensitivity of the radioligand for changes in cardiac expression induced by DOX. Moreover, its high basal expression in human and murine heart **(Figure S9)**, may render TSPO imaging insensitive to small changes in expression levels. Additionally, the systemic inflammation induced by DOX treatment in this mouse model may further obscure small changes in cardiac PET signal and may represent a limitation of our model.

Our decision to target NET with [^18^F]MFBG was based on prior evidence that cardiac uptake of radiolabeled *meta*-iodobenzylguanidine (MIBG) decreases in a dose-dependent manner in rodents treated with anthracyclines^57^ and in cancer patients that had received anthracycline chemotherapy relative to those receiving alternative treatment.^38,105^ We saw considerable cardiac uptake normalized to skeletal muscle of [^18^F]MFBG in both DOX animals and CTRL (**Figure S3**), but no decline over the 12 W observation period. By contrast, in the early stages of cardiotoxicity, [^18^F]MFBG uptake was actually higher in DOX mice, though this was not statistically significant (1 W; *p* = 0.400, 2 W; *p* = 0.527, and 4 W; *p* = 0.161). Although prior studies did show declines in [^123/125^I]MIBG uptake concurrent with LVEF decline, the uptake deficit was sustained. It may be that our follow up period was too short to detect differences between our groups, but the convergence of the curves in **Figure S3** suggests that differences are unlikely to emerge. A retrospective analysis could not discriminate asymptomatic pediatric cancer survivors from healthy controls using [^123^I]MIBG image quantification, and myocardial sympathetic activity was neither related to anthracycline dose nor LVEF.^106^ This may indicate that sympathetic denervation is not sufficiently pronounced in chronic cardiotoxicity to represent a reliable imaging biomarker.

To date, echo has been used in cardio-oncology as the main imaging modality for screening patients with suspected cardiotoxicity.^107^ Given the implementation of new echo techniques that improve its sensitivity for subclinical disease and the continued definition of cardiotoxicity in terms of LVEF decreases, echo will continue to play a major role in diagnosis and monitoring progression. However, our results support a role for PET imaging in the management of cancer patients receiving anthracycline chemotherapy. In our model, we detected pathological cardiac remodeling in DOX hearts as much as 8 weeks before functional decline was evident by echo. Early diagnosis of cardiotoxicity could greatly improve its treatment, as evidenced by the more complete recovery of LVEF in patients with cardiotoxicity administered ACE inhibitors or beta blockers shortly after anthracycline chemotherapy than patients treated a few months later.^108^ Molecular imaging techniques such as FAPI PET may lead to even more impressive treatment outcomes by identifying the activation of specific pathological pathways whose inhibition could mitigate or even prevent cardiotoxicity. For example FAP inhibition improves cardiac repair after myocardial infarction.^109,110^ Future work is needed determine whether it is similarly beneficial in cardiotoxicity, but this example does highlight the potential benefit of PET imaging biomarkers in treating cardiotoxicity.

We acknowledge several limitations of our study. Firstly, although we showed the correlation between cardiac FAPI PET uptake and FAP expression, elevated background FAPI PET signal in DOX mice was also seen due to uptake in the gastrointestinal region, muscle, and in some cases, lung. This likely reflects off-target uptake due to sustained and global inflammation caused by systemic administration of DOX. This phenomenon was previously observed in FAPI PET imaging of a pre-clinical model of idiopathic pulmonary fibrosis induced by bleomycin.^111^ Secondly, our methods of quantifying FAP protein expression could not distinguish between membrane-bound FAP and cytoplasmic FAP. As our radioligand does not cross the cell membrane, the signals derived from this probe reflect the binding of membrane-bound FAP. To our knowledge, FAP is primarily an outer membrane protein, though increased cytoplasmic expression was recently reported in lung adenocarcinoma cells.^112^ We therefore cannot rule out the possibility that cytoplasmic FAP protein expression confounds our analysis even though our results identify increased FAP protein expression and activity and gene expression in the DOX mice. Moreover, our studies have not determined the function of FAP in DOX-induced cardiotoxicity. Contrary to our expectations, increased FAP expression did not result in increased fibrosis. We speculate that FAP is broadly involved in ECM remodeling, but without identifying its specific role in this pathology, it will be challenging to determine whether FAPI PET could also be a prognostic biomarker in cardiotoxicity. Larger sample sizes could possibly determine whether early increases in FAPI PET correspond to larger declines in functional parameters such as LVEF. Finally, our studies were limited to male mice because female mice are less susceptible to cardiotoxicity.^11^ Therefore, further research is required to determine whether FAPI PET will be equally valuable in female patients.

## CONCLUSIONS

Although anthracycline chemotherapy has dramatically improved treatment outcome in cancer patients, especially in children with cancer, it causes cardiotoxicity with an increased risk of heart failure in a significant number of patients. Existing imaging modalities detect cardiac functional deficits but do not identify the underlying, potentially treatable, pathologies responsible for these deficits. We demonstrate a significant and sustained increase of FAP expression in response to systemic administration of doxorubicin and show that this change can be imaged by PET using [^68^Ga]Ga-FAPI-04. Functional changes were not evident by routine echo until 10 weeks, as much as 8 weeks after cardiac FAPI PET signal increases were detected. These findings suggest that FAPI PET is a diagnostic imaging biomarker for incipient cardiotoxicity and a potential complement to echo for the management of cancer patients receiving anthracycline chemotherapy. Early detection of FAP-mediated cardiac remodeling may improve the efficacy of therapeutic interventions to delay or even prevent heart failure.

## ACKNOWLEDGMENTS

The authors wish to acknowledge Anil Ekkati, Ph.D. and David J. Warren, Ph.D. of the Milstein Core Chemistry facility for synthesizing DPA-714 and its tosylate precursor, and Pradeep K. Singh, Ph.D., and Stephen G. DiMagno, Ph.D. of the University of Illinois-Chicago for support with ALP-mFBG synthesis. The authors also wish to acknowledge Alejandro Amor-Coarasa, Ph.D., of Ratio Therapeutics, Serge Lyashchenko, Pharm.D., and Eva Burnazi of Memorial-Sloan Kettering Cancer Center for helpful discussions concerning the radiosynthesis of [^18^F]MFBG.

## AUTHOR CONTRIBUTIONS

The studies were conceived by J.M.K., J.W.B., and A.diL. Experiments were designed by C-H.L. and J.M.K. and performed by C-H.L, O.M., L.R., and T.M.J. Data analysis was performed by C-H.L., O.M., S. Cho., T.M.J., A. diL., and J.M.K. Pathology and quantitation of ISH and IHC was performed by S.E.C. Funding was acquired by J.M.K. The manuscript was written by C-H.L. and J.M.K. and reviewed by O.M., L.R., S.E.C., S. Cho., T.M.J., J.W.B., and A. diL.

## Nonstandard Abbreviations and Acronyms

ACE: Acetylcholinesterase
BP: Biological process
BSA: Bovine serum albumin
CT: Computed tomography
CTRL: Control
DEG: Differentially expressed genes
DOX: Doxorubicin
FAP: Fibroblast activation protein alpha
FAPI: Fibroblast activation protein alpha inhibitor
FC: Fold change
FPKM: Fragments per kilobase of transcript per million mapped reads FS Fractional shortening
GO: Gene oncology
HW: Heart weight
ID: Injected dose
ISH: In situ hybridization
LVDd: Left ventricle end-diastolic diameter
LVDs: Left ventricle end-systolic diameter
LVEF: Left ventricular ejection fraction
MFBG: meta-Fluorobenzylguanidine
MIBG: meta-Iodobenzylguanidine
MIP: Maximum intensity projection
NET: Norepinephrine transporter
PBR: Peripheral-type benzodiazepine receptor
PBS: Phosphate buffered saline
PET: Positron emission tomography
RNA-seq: NA sequencing
ROI: Region of interest
SPECT: Single photon computed tomography TL Tibia length
Top2β: Topoisomerase-2β
TSPO: Translocator protein 18-kDa
VOI: Volume of interest
WB: Western blot

## REFERENCES

1. Asselin BL, Devidas M, Chen L, Franco VI, Pullen J, Borowitz MJ, Hutchison RE, Ravindranath Y, Armenian SH, Camitta BM, Lipshultz SE. Cardioprotection and Safety of Dexrazoxane in Patients Treated for Newly Diagnosed T-Cell Acute Lymphoblastic Leukemia or Advanced-Stage Lymphoblastic Non-Hodgkin Lymphoma: A Report of the Children’s Oncology Group Randomized Trial Pediatric Oncology Group 9404. J Clin Oncol. 2016;34:854–862. doi: 10.1200/JCO.2015.60.8851

2. McGowan JV, Chung R, Maulik A, Piotrowska I, Walker JM, Yellon DM. Anthracycline Chemotherapy and Cardiotoxicity. Cardiovasc Drugs Ther. 2017;31:63–75. doi: 10.1007/s10557-016-6711-0

3. van Dalen EC, Raphael MF, Caron HN, Kremer LC. Treatment including anthracyclines versus treatment not including anthracyclines for childhood cancer. Cochrane Database Syst Rev. 2014:CD006647. doi: 10.1002/14651858.CD006647.pub4

4. Unverferth BJ, Magorien RD, Balcerzak SP, Leier CV, Unverferth DV. Early changes in human myocardial nuclei after doxorubicin. Cancer. 1983;52:215–221. doi: 10.1002/1097-0142(19830715)52:2<215::aid-cncr2820520206>3.0.co;2-f

5. Sawyer DB. Anthracyclines and heart failure. N Engl J Med. 2013;368:1154–1156. doi: 10.1056/NEJMcibr1214975

6. Von Hoff DD, Layard MW, Basa P, Davis HL, Jr., Von Hoff AL, Rozencweig M, Muggia FM. Risk factors for doxorubicin-induced congestive heart failure. Ann Intern Med. 1979;91:710–717. doi: 10.7326/0003-4819-91-5-710

7. Totzeck M, Siebermair J, Rassaf T, Rischpler C. Cardiac fibroblast activation detected by positron emission tomography/computed tomography as a possible sign of cardiotoxicity. Eur Heart J. 2020;41:1060. doi: 10.1093/eurheartj/ehz736

8. Mancilla TR, Iskra B, Aune GJ. Doxorubicin-Induced Cardiomyopathy in Children. Compr Physiol. 2019;9:905–931. doi: 10.1002/cphy.c180017

9. Harake D, Franco VI, Henkel JM, Miller TL, Lipshultz SE. Cardiotoxicity in childhood cancer survivors: strategies for prevention and management. Future Cardiol. 2012;8:647–670. doi: 10.2217/fca.12.44

10. Cardinale D, Iacopo F, Cipolla CM. Cardiotoxicity of Anthracyclines. Front Cardiovasc Med. 2020;7:26. doi: 10.3389/fcvm.2020.00026

11. Willis MS, Parry TL, Brown DI, Mota RI, Huang W, Beak JY, Sola M, Zhou C, Hicks ST, Caughey MC, et al. Doxorubicin Exposure Causes Subacute Cardiac Atrophy Dependent on the Striated Muscle-Specific Ubiquitin Ligase MuRF1. Circ Heart Fail. 2019;12:e005234. doi: 10.1161/CIRCHEARTFAILURE.118.005234

12. Sawaya H, Sebag IA, Plana JC, Januzzi JL, Ky B, Tan TC, Cohen V, Banchs J, Carver JR, Wiegers SE, et al. Assessment of echocardiography and biomarkers for the extended prediction of cardiotoxicity in patients treated with anthracyclines, taxanes, and trastuzumab. Circ Cardiovasc Imaging. 2012;5:596–603. doi: 10.1161/CIRCIMAGING.112.973321

13. Jenkins C, Bricknell K, Hanekom L, Marwick TH. Reproducibility and accuracy of echocardiographic measurements of left ventricular parameters using real-time three-dimensional echocardiography. J Am Coll Cardiol. 2004;44:878–886. doi: 10.1016/j.jacc.2004.05.050

14. Hoffmann R, von Bardeleben S, ten Cate F, Borges AC, Kasprzak J, Firschke C, Lafitte S, Al-Saadi N, Kuntz-Hehner S, Engelhardt M, et al. Assessment of systolic left ventricular function: a multi-centre comparison of cineventriculography, cardiac magnetic resonance imaging, unenhanced and contrast-enhanced echocardiography. Eur Heart J. 2005;26:607–616. doi: 10.1093/eurheartj/ehi083

15. Thavendiranathan P, Grant AD, Negishi T, Plana JC, Popovic ZB, Marwick TH. Reproducibility of echocardiographic techniques for sequential assessment of left ventricular ejection fraction and volumes: application to patients undergoing cancer chemotherapy. J Am Coll Cardiol. 2013;61:77–84. doi: 10.1016/j.jacc.2012.09.035

16. Ky B, Putt M, Sawaya H, French B, Januzzi JL, Jr., Sebag IA, Plana JC, Cohen V, Banchs J, Carver JR, et al. Early increases in multiple biomarkers predict subsequent cardiotoxicity in patients with breast cancer treated with doxorubicin, taxanes, and trastuzumab. J Am Coll Cardiol. 2014;63:809–816. doi: 10.1016/j.jacc.2013.10.061

17. Becker MMC, Arruda GFA, Berenguer DRF, Buril RO, Cardinale D, Brandao SCS. Anthracycline cardiotoxicity: current methods of diagnosis and possible role of (18)F-FDG PET/CT as a new biomarker. Cardiooncology. 2023;9:17. doi: 10.1186/s40959-023-00161-6

18. Liu J, Banchs J, Mousavi N, Plana JC, Scherrer-Crosbie M, Thavendiranathan P, Barac A. Contemporary Role of Echocardiography for Clinical Decision Making in Patients During and After Cancer Therapy. JACC Cardiovasc Imaging. 2018;11:1122–1131. doi: 10.1016/j.jcmg.2018.03.025

19. Jordan JH, Todd RM, Vasu S, Hundley WG. Cardiovascular Magnetic Resonance in the Oncology Patient. JACC Cardiovasc Imaging. 2018;11:1150–1172. doi: 10.1016/j.jcmg.2018.06.004

20. Plana JC, Thavendiranathan P, Bucciarelli-Ducci C, Lancellotti P. Multi-Modality Imaging in the Assessment of Cardiovascular Toxicity in the Cancer Patient. JACC Cardiovasc Imaging. 2018;11:1173–1186. doi: 10.1016/j.jcmg.2018.06.003

21. Yu AF, Ky B. Roadmap for biomarkers of cancer therapy cardiotoxicity. Heart. 2016;102:425–430. doi: 10.1136/heartjnl-2015-307894

22. Polomski ES, Antoni ML, Jukema JW, Kroep JR, Dibbets-Schneider P, Sattler MGA, de Geus-Oei LF. Nuclear medicine imaging methods of radiation-induced cardiotoxicity. Semin Nucl Med. 2022;52:597–610. doi: 10.1053/j.semnuclmed.2022.02.001

23. Vogelsang TW, Jensen RJ, Hesse B, Kjaer A. BNP cannot replace gated equilibrium radionuclide ventriculography in monitoring of anthracycline-induced cardiotoxity. Int J Cardiol. 2008;124:193–197. doi: 10.1016/j.ijcard.2007.02.003

24. Zeiss CJ, Gatti DM, Toro-Salazar O, Davis C, Lutz CM, Spinale F, Stearns T, Furtado MB, Churchill GA. Doxorubicin-Induced Cardiotoxicity in Collaborative Cross (CC) Mice Recapitulates Individual Cardiotoxicity in Humans. G3 (Bethesda). 2019;9:2637–2646. doi: 10.1534/g3.119.400232

25. Moses WW. Fundamental Limits of Spatial Resolution in PET. Nucl Instrum Methods Phys Res A. 2011;648 Supplement 1:S236–S240. doi: 10.1016/j.nima.2010.11.092

26. Loktev A, Lindner T, Mier W, Debus J, Altmann A, Jager D, Giesel F, Kratochwil C, Barthe P, Roumestand C, Haberkorn U. A Tumor-Imaging Method Targeting Cancer-Associated Fibroblasts. J Nucl Med. 2018;59:1423–1429. doi: 10.2967/jnumed.118.210435

27. Lindner T, Loktev A, Altmann A, Giesel F, Kratochwil C, Debus J, Jager D, Mier W, Haberkorn U. Development of Quinoline-Based Theranostic Ligands for the Targeting of Fibroblast Activation Protein. J Nucl Med. 2018;59:1415–1422. doi: 10.2967/jnumed.118.210443

28. Loktev A, Lindner T, Burger EM, Altmann A, Giesel F, Kratochwil C, Debus J, Marme F, Jager D, Mier W, Haberkorn U. Development of Fibroblast Activation Protein-Targeted Radiotracers with Improved Tumor Retention. J Nucl Med. 2019;60:1421–1429. doi: 10.2967/jnumed.118.224469

29. Lindner T, Altmann A, Giesel F, Kratochwil C, Kleist C, Kramer S, Mier W, Cardinale J, Kauczor HU, Jager D, et al. (18)F-labeled tracers targeting fibroblast activation protein. EJNMMI Radiopharm Chem. 2021;6:26. doi: 10.1186/s41181-021-00144-x

30. James ML, Fulton RR, Vercoullie J, Henderson DJ, Garreau L, Chalon S, Dolle F, Costa B, Guilloteau D, Kassiou M. DPA-714, a new translocator protein-specific ligand: synthesis, radiofluorination, and pharmacologic characterization. J Nucl Med. 2008;49:814–822. doi: 10.2967/jnumed.107.046151

31. MacAskill MG, Stadulyte A, Williams L, Morgan TEF, Sloan NL, Alcaide-Corral CJ, Walton T, Wimberley C, McKenzie CA, Spath N, et al. Quantification of Macrophage-Driven Inflammation During Myocardial Infarction with (18)F-LW223, a Novel TSPO Radiotracer with Binding Independent of the rs6971 Human Polymorphism. J Nucl Med. 2021;62:536–544. doi: 10.2967/jnumed.120.243600

32. Simeon FG, Lee JH, Morse CL, Stukes I, Zoghbi SS, Manly LS, Liow JS, Gladding RL, Dick RM, Yan X, et al. Synthesis and Screening in Mice of Fluorine-Containing PET Radioligands for TSPO: Discovery of a Promising (18)F-Labeled Ligand. J Med Chem. 2021;64:16731–16745. doi: 10.1021/acs.jmedchem.1c01562

33. Pandit-Taskar N, Zanzonico P, Staton KD, Carrasquillo JA, Reidy-Lagunes D, Lyashchenko S, Burnazi E, Zhang H, Lewis JS, Blasberg R, et al. Biodistribution and Dosimetry of (18)F-Meta-Fluorobenzylguanidine: A First-in-Human PET/CT Imaging Study of Patients with Neuroendocrine Malignancies. J Nucl Med. 2018;59:147–153. doi: 10.2967/jnumed.117.193169

34. Yu M, Bozek J, Lamoy M, Guaraldi M, Silva P, Kagan M, Yalamanchili P, Onthank D, Mistry M, Lazewatsky J, et al. Evaluation of LMI1195, a novel 18F-labeled cardiac neuronal PET imaging agent, in cells and animal models. Circ Cardiovasc Imaging. 2011;4:435–443. doi: 10.1161/CIRCIMAGING.110.962126

35. Arlicot N, Vercouillie J, Ribeiro MJ, Tauber C, Venel Y, Baulieu JL, Maia S, Corcia P, Stabin MG, Reynolds A, et al. Initial evaluation in healthy humans of [18F]DPA-714, a potential PET biomarker for neuroinflammation. Nucl Med Biol. 2012;39:570–578. doi: 10.1016/j.nucmedbio.2011.10.012

36. Lencova-Popelova O, Jirkovsky E, Mazurova Y, Lenco J, Adamcova M, Simunek T, Gersl V, Sterba M. Molecular remodeling of left and right ventricular myocardium in chronic anthracycline cardiotoxicity and post-treatment follow up. PLoS One. 2014;9:e96055. doi: 10.1371/journal.pone.0096055

37. Fabiani I, Aimo A, Grigoratos C, Castiglione V, Gentile F, Saccaro LF, Arzilli C, Cardinale D, Passino C, Emdin M. Oxidative stress and inflammation: determinants of anthracycline cardiotoxicity and possible therapeutic targets. Heart Fail Rev. 2021;26:881–890. doi: 10.1007/s10741-020-10063-9

38. Valdes Olmos RA, Ten Bokkel Huinink WW, Tenhoeve RFA, Vantinteren H, Bruning PF, Vanvlies B, Hoefnagel CA. Assessment of Anthracycline-Related Myocardial Adrenergic Derangement by [I-123] Metaiodobenzylguanidine Scintigraphy. Eur J Cancer. 1995;31a:26–31. doi: Doi 10.1016/0959-8049(94)00357-B

39. Amgalan D, Garner TP, Pekson R, Jia XF, Yanamandala M, Paulino V, Liang FG, Corbalan JJ, Lee J, Chen Y, et al. A small-molecule allosteric inhibitor of BAX protects against doxorubicin-induced cardiomyopathy. Nat Cancer. 2020;1:315–328. doi: 10.1038/s43018-020-0039-1

40. Sasset L, Manzo OL, Zhang Y, Marino A, Rubinelli L, Riemma MA, Chalasani MLS, Dasoveanu DC, Roviezzo F, Jankauskas SS, et al. Nogo-A reduces ceramide de novo biosynthesis to protect from heart failure. Cardiovasc Res. 2023;119:506–519. doi: 10.1093/cvr/cvac108

41. Mitchell C, Rahko PS, Blauwet LA, Canaday B, Finstuen JA, Foster MC, Horton K, Ogunyankin KO, Palma RA, Velazquez EJ. Guidelines for Performing a Comprehensive Transthoracic Echocardiographic Examination in Adults: Recommendations from the American Society of Echocardiography. J Am Soc Echocardiogr. 2019;32:1–64. doi: 10.1016/j.echo.2018.06.004

42. Loening AM, Gambhir SS. AMIDE: a free software tool for multimodality medical image analysis. Mol Imaging. 2003;2:131–137. doi: 10.1162/15353500200303133

43. Lee CH, Kim MJ, Lee HH, Paeng JC, Park YJ, Oh SW, Chai YJ, Kim YA, Cheon GJ, Kang KW, et al. Adenine Nucleotide Translocase 2 as an Enzyme Related to [(18)F] FDG Accumulation in Various Cancers. Mol Imaging Biol. 2019;21:722–730. doi: 10.1007/s11307-018-1268-x

44. Dobin A, Davis CA, Schlesinger F, Drenkow J, Zaleski C, Jha S, Batut P, Chaisson M, Gingeras TR. STAR: ultrafast universal RNA-seq aligner. Bioinformatics. 2013;29:15–21. doi: 10.1093/bioinformatics/bts635

45. Trapnell C, Williams BA, Pertea G, Mortazavi A, Kwan G, van Baren MJ, Salzberg SL, Wold BJ, Pachter L. Transcript assembly and quantification by RNA-Seq reveals unannotated transcripts and isoform switching during cell differentiation. Nat Biotechnol. 2010;28:511–515. doi: 10.1038/nbt.1621

46. Trapnell C, Hendrickson DG, Sauvageau M, Goff L, Rinn JL, Pachter L. Differential analysis of gene regulation at transcript resolution with RNA-seq. Nat Biotechnol. 2013;31:46–53. doi: 10.1038/nbt.2450

47. Anders S, Pyl PT, Huber W. HTSeq--a Python framework to work with high-throughput sequencing data. Bioinformatics. 2015;31:166–169. doi: 10.1093/bioinformatics/btu638

48. Love MI, Huber W, Anders S. Moderated estimation of fold change and dispersion for RNA-seq data with DESeq2. Genome Biol. 2014;15:550. doi: 10.1186/s13059-014-0550-8

49. von Mering C, Huynen M, Jaeggi D, Schmidt S, Bork P, Snel B. STRING: a database of predicted functional associations between proteins. Nucleic Acids Res. 2003;31:258–261. doi: 10.1093/nar/gkg034

50. Zhang S, Liu X, Bawa-Khalfe T, Lu LS, Lyu YL, Liu LF, Yeh ET. Identification of the molecular basis of doxorubicin-induced cardiotoxicity. Nat Med. 2012;18:1639–1642. doi: 10.1038/nm.2919

51. Ruiz-Villalba A, Mattiotti A, Gunst QD, Cano-Ballesteros S, van den Hoff MJ, Ruijter JM. Reference genes for gene expression studies in the mouse heart. Sci Rep. 2017;7:24. doi: 10.1038/s41598-017-00043-9

52. Goldsmith EC, Bradshaw AD, Zile MR, Spinale FG. Myocardial fibroblast-matrix interactions and potential therapeutic targets. J Mol Cell Cardiol. 2014;70:92–99. doi: 10.1016/j.yjmcc.2014.01.008

53. Moita MR, Silva MM, Diniz C, Serra M, Hoet RM, Barbas A, Simao D. Transcriptome and proteome profiling of activated cardiac fibroblasts supports target prioritization in cardiac fibrosis. Front Cardiovasc Med. 2022;9:1015473. doi: 10.3389/fcvm.2022.1015473

54. Kelly JM, Babich JW. PET Tracers for Imaging Cardiac Function in Cardio-oncology. Curr Cardiol Rep. 2022;24:247–260. doi: 10.1007/s11886-022-01641-4

55. Safee ZM, Baark F, Waters ECT, Veronese M, Pell VR, Clark JE, Mota F, Livieratos L, Eykyn TR, Blower PJ, Southworth R. Detection of anthracycline-induced cardiotoxicity using perfusion-corrected (99m)Tc sestamibi SPECT. Sci Rep. 2019;9:216. doi: 10.1038/s41598-018-36721-5

56. McCluskey SP, Haslop A, Coello C, Gunn RN, Tate EW, Southworth R, Plisson C, Long NJ, Wells LA. Imaging of Chemotherapy-Induced Acute Cardiotoxicity with (18)F-Labeled Lipophilic Cations. J Nucl Med. 2019;60:1750–1756. doi: 10.2967/jnumed.119.226787

57. Wakasugi S, Fischman AJ, Babich JW, Aretz HT, Callahan RJ, Nakaki M, Wilkinson R, Strauss HW. Metaiodobenzylguanidine: evaluation of its potential as a tracer for monitoring doxorubicin cardiomyopathy. J Nucl Med. 1993;34:1283–1286.

58. Collin B, Oudot A, Vrigneaud JM, Delemasure S, Guillemin M, Walker PM, Lalande A, Cochet A, Brunotte F. Abnormal cardiac adrenergic neuron activity assessed by I-123-MIBG is an early marker of cardiac dysfunction in doxorubicin-induced cardiomyopathy in rats. Eur J Nucl Med Mol I. 2016;43:S88–S89.

59. Hartmann F, Ziegler S, Nekolla S, Hadamitzky M, Seyfarth M, Richardt G, Schwaiger M. Regional patterns of myocardial sympathetic denervation in dilated cardiomyopathy: an analysis using carbon-11 hydroxyephedrine and positron emission tomography. Heart. 1999;81:262–270. doi: 10.1136/hrt.81.3.262

60. Thackeray JT, Hupe HC, Wang Y, Bankstahl JP, Berding G, Ross TL, Bauersachs J, Wollert KC, Bengel FM. Myocardial Inflammation Predicts Remodeling and Neuroinflammation After Myocardial Infarction. J Am Coll Cardiol. 2018;71:263–275. doi: 10.1016/j.jacc.2017.11.024

61. Borchert T, Hess A, Lukacevic M, Ross TL, Bengel FM, Thackeray JT. Angiotensin-converting enzyme inhibitor treatment early after myocardial infarction attenuates acute cardiac and neuroinflammation without effect on chronic neuroinflammation. Eur J Nucl Med Mol Imaging. 2020;47:1757–1768. doi: 10.1007/s00259-020-04736-8

62. Glasenapp A, Derlin K, Gutberlet M, Hess A, Ross TL, Wester HJ, Bengel FM, Thackeray JT. Molecular Imaging of Inflammation and Fibrosis in Pressure Overload Heart Failure. Circ Res. 2021;129:369–382. doi: 10.1161/Circresaha.120.318539

63. Jenkins WS, Vesey AT, Stirrat C, Connell M, Lucatelli C, Neale A, Moles C, Vickers A, Fletcher A, Pawade T, et al. Cardiac alpha(V)beta(3) integrin expression following acute myocardial infarction in humans. Heart. 2017;103:607–615. doi: 10.1136/heartjnl-2016-310115

64. Varasteh Z, Mohanta S, Robu S, Braeuer M, Li Y, Omidvari N, Topping G, Sun T, Nekolla SG, Richter A, et al. Molecular Imaging of Fibroblast Activity After Myocardial Infarction Using a (68)Ga-Labeled Fibroblast Activation Protein Inhibitor, FAPI-04. J Nucl Med. 2019;60:1743–1749. doi: 10.2967/jnumed.119.226993

65. Heckmann MB, Reinhardt F, Finke D, Katus HA, Haberkorn U, Leuschner F, Lehmann LH. Relationship Between Cardiac Fibroblast Activation Protein Activity by Positron Emission Tomography and Cardiovascular Disease. Circ-Cardiovasc Imag. 2020;13. doi: ARTN e010628. 10.1161/CIRCIMAGING.120.010628

66. Siebermair J, Kohler MI, Kupusovic J, Nekolla SG, Kessler L, Ferdinandus J, Guberina N, Stuschke M, Grafe H, Siveke JT, et al. Cardiac fibroblast activation detected by Ga-68 FAPI PET imaging as a potential novel biomarker of cardiac injury/remodeling. J Nucl Cardiol. 2021;28:812–821. doi: 10.1007/s12350-020-02307-w

67. Langer LBN, Hess A, Korkmaz Z, Tillmanns J, Reffert LM, Bankstahl JP, Bengel FM, Thackeray JT, Ross TL. Molecular imaging of fibroblast activation protein after myocardial infarction using the novel radiotracer [Ga-68]MHLL1. Theranostics. 2021;11:7755–7766. doi: 10.7150/thno.51419

68. O’Farrell AC, Evans R, Silvola JM, Miller IS, Conroy E, Hector S, Cary M, Murray DW, Jarzabek MA, Maratha A, et al. A Novel Positron Emission Tomography (PET) Approach to Monitor Cardiac Metabolic Pathway Remodeling in Response to Sunitinib Malate. PLoS One. 2017;12:e0169964. doi: 10.1371/journal.pone.0169964

69. Croteau E, Tremblay S, Gascon S, Dumulon-Perreault V, Labbe SM, Rousseau JA, Cunnane SC, Carpentier AC, Benard F, Lecomte R. [(11)C]-Acetoacetate PET imaging: a potential early marker for cardiac heart failure. Nucl Med Biol. 2014;41:863–870. doi: 10.1016/j.nucmedbio.2014.08.006

70. Sarocchi M, Bauckneht M, Arboscello E, Capitanio S, Marini C, Morbelli S, Miglino M, Congiu AG, Ghigliotti G, Balbi M, et al. An increase in myocardial 18-fluorodeoxyglucose uptake is associated with left ventricular ejection fraction decline in Hodgkin lymphoma patients treated with anthracycline. J Transl Med. 2018;16:295. doi: 10.1186/s12967-018-1670-9

71. Kratochwil C, Flechsig P, Lindner T, Abderrahim L, Altmann A, Mier W, Adeberg S, Rathke H, Rohrich M, Winter H, et al. (68)Ga-FAPI PET/CT: Tracer Uptake in 28 Different Kinds of Cancer. J Nucl Med. 2019;60:801–805. doi: 10.2967/jnumed.119.227967

72. Mori Y, Dendl K, Cardinale J, Kratochwil C, Giesel FL, Haberkorn U. FAPI PET: Fibroblast Activation Protein Inhibitor Use in Oncologic and Nononcologic Disease. Radiology. 2023;306:e220749. doi: 10.1148/radiol.220749

73. Peyronneau MA, Kuhnast B, Nguyen DL, Jego B, Sayet G, Caille F, Lavisse S, Gervais P, Stankoff B, Sarazin M, et al. [(18)F]DPA-714: Effect of co-medications, age, sex, BMI and TSPO polymorphism on the human plasma input function. Eur J Nucl Med Mol Imaging. 2023. doi: 10.1007/s00259-023-06286-1

74. Pauwels E, Celen S, Vandamme M, Leysen W, Baete K, Bechter O, Bex M, Serdons K, Van Laere K, Bormans G, Deroose CM. Improved resolution and sensitivity of [(18)F]MFBG PET compared with [(123)I]MIBG SPECT in a patient with a norepinephrine transporter-expressing tumour. Eur J Nucl Med Mol Imaging. 2021;48:313–315. doi: 10.1007/s00259-020-04830-x

75. He X, Du T, Long T, Liao X, Dong Y, Huang ZP. Signaling cascades in the failing heart and emerging therapeutic strategies. Signal Transduct Target Ther. 2022;7:134. doi: 10.1038/s41392-022-00972-6

76. Capranico G, Tinelli S, Austin CA, Fisher ML, Zunino F. Different patterns of gene expression of topoisomerase II isoforms in differentiated tissues during murine development. Biochim Biophys Acta. 1992;1132:43–48. doi: 10.1016/0167-4781(92)90050-a

77. Vejpongsa P, Yeh ET. Topoisomerase 2beta: a promising molecular target for primary prevention of anthracycline-induced cardiotoxicity. Clin Pharmacol Ther. 2014;95:45–52. doi: 10.1038/clpt.2013.201

78. Timm KN, Perera C, Ball V, Henry JA, Miller JJ, Kerr M, West JA, Sharma E, Broxholme J, Logan A, et al. Early detection of doxorubicin-induced cardiotoxicity in rats by its cardiac metabolic signature assessed with hyperpolarized MRI. Commun Biol. 2020;3:692. doi: 10.1038/s42003-020-01440-z

79. Sadoshima J, Izumo S. The cellular and molecular response of cardiac myocytes to mechanical stress. Annu Rev Physiol. 1997;59:551–571. doi: 10.1146/annurev.physiol.59.1.551

80. Kehat I, Molkentin JD. Molecular pathways underlying cardiac remodeling during pathophysiological stimulation. Circulation. 2010;122:2727–2735. doi: 10.1161/CIRCULATIONAHA.110.942268

81. Wang H, Wu Q, Liu Z, Luo X, Fan Y, Liu Y, Zhang Y, Hua S, Fu Q, Zhao M, et al. Downregulation of FAP suppresses cell proliferation and metastasis through PTEN/PI3K/AKT and Ras-ERK signaling in oral squamous cell carcinoma. Cell Death Dis. 2014;5:e1155. doi: 10.1038/cddis.2014.122

82. Jia J, Martin TA, Ye L, Jiang WG. FAP-alpha (Fibroblast activation protein-alpha) is involved in the control of human breast cancer cell line growth and motility via the FAK pathway. Bmc Cell Biol. 2014;15. doi: Artn 16. 10.1186/1471-2121-15-16

83. Yang XG, Lin YL, Shi YH, Li BJ, Liu WR, Yin W, Dang YJ, Chu YW, Fan J, He R. FAP Promotes Immunosuppression by Cancer-Associated Fibroblasts in the Tumor Microenvironment via STAT3-CCL2 Signaling. Cancer Res. 2016;76:4124–4135. doi: 10.1158/0008-5472.Can-15-2973

84. Wang XM, Yu DMT, McCaughan GW, Gorrell MD. Fibroblast activation protein increases apoptosis, cell adhesion, and migration by the LX-2 human stellate cell line. Hepatology. 2005;42:935–945. doi: 10.1002/hep.20853

85. Chen M, Lei X, Shi C, Huang M, Li X, Wu B, Li Z, Han W, Du B, Hu J, et al. Pericyte-targeting prodrug overcomes tumor resistance to vascular disrupting agents. J Clin Invest. 2017;127:3689–3701. doi: 10.1172/JCI94258

86. Jellis C, Martin J, Narula J, Marwick TH. Assessment of Nonischemic Myocardial Fibrosis. Journal of the American College of Cardiology. 2010;56:89–97. doi: 10.1016/j.jacc.2010.02.047

87. Ramirez-Montagut T, Blachere NE, Sviderskaya EV, Bennett DC, Rettig WJ, Garin-Chesa P, Houghton AN. FAP alpha, a surface peptidase expressed during wound healing, is a tumor suppressor. Oncogene. 2004;23:5435–5446. doi: 10.1038/sj.onc.1207730

88. Travers JG, Kamal FA, Robbins J, Yutzey KE, Blaxall BC. Cardiac Fibrosis The Fibroblast Awakens. Circ Res. 2016;118:1021–1040. doi: 10.1161/Circresaha.115.306565

89. Tillmanns J, Hoffmann D, Habbaba Y, Schmitto JD, Sedding D, Fraccarollo D, Galuppo P, Bauersachs J. Fibroblast activation protein alpha expression identifies activated fibroblasts after myocardial infarction. Journal of Molecular and Cellular Cardiology. 2015;87:194–203. doi: 10.1016/j.yjmcc.2015.08.016

90. Shi X, Lin X, Huo L, Li X. Cardiac fibroblast activation in dilated cardiomyopathy detected by positron emission tomography. J Nucl Cardiol. 2022;29:881–884. doi: 10.1007/s12350-020-02315-w

91. Wang L, Zhang Z, Zhao Z, Yan C, Fang W. (68)Ga-FAPI right heart uptake in a patient with idiopathic pulmonary arterial hypertension. J Nucl Cardiol. 2022;29:1475–1477. doi: 10.1007/s12350-020-02407-7

92. Song WY, Zhang X, He SK, Gai YK, Qin CX, Hu F, Wang Y, Wang ZH, Bai P, Wang J, Lan XL. Ga-68-FAPI PET visualize heart failure: from mechanism to clinic. Eur J Nucl Med Mol I. 2023;50:475–485. doi: 10.1007/s00259-022-05994-4

93. Wu C, Macleod I, Su AI. BioGPS and MyGene.info: organizing online, gene-centric information. Nucleic Acids Res. 2013;41:D561–565. doi: 10.1093/nar/gks1114

94. Jordan JH, Castellino SM, Melendez GC, Klepin HD, Ellis LR, Lamar Z, Vasu S, Kitzman DW, Ntim WO, Brubaker PH, et al. Left Ventricular Mass Change After Anthracycline Chemotherapy. Circ Heart Fail. 2018;11:e004560. doi: 10.1161/CIRCHEARTFAILURE.117.004560

95. Harvey PA, Leinwand LA. Cardiac atrophy and remodeling. In: Cellular and molecular pathobiology of cardiovascular disease. Elsevier; 2014:37–50.

96. Nishi M, Wang PY, Hwang PM. Cardiotoxicity of Cancer Treatments: Focus on Anthracycline Cardiomyopathy. Arterioscler Thromb Vasc Biol. 2021;41:2648–2660. doi: 10.1161/ATVBAHA.121.316697

97. Thackeray JT. Molecular Imaging Using Cardiac PET/CT: Opportunities to Harmonize Diagnosis and Therapy. Curr Cardiol Rep. 2021;23:96. doi: 10.1007/s11886-021-01526-y

98. Morin D, Musman J, Pons S, Berdeaux A, Ghaleh B. Mitochondrial translocator protein (TSPO): From physiology to cardioprotection. Biochem Pharmacol. 2016;105:1–13. doi: 10.1016/j.bcp.2015.12.003

99. Largeau B, Dupont AC, Guilloteau D, Santiago-Ribeiro MJ, Arlicot N. TSPO PET Imaging: From Microglial Activation to Peripheral Sterile Inflammatory Diseases? Contrast Media Mol Imaging. 2017;2017:6592139. doi: 10.1155/2017/6592139

100. Narayan N, Owen DR, Mandhair H, Smyth E, Carlucci F, Saleem A, Gunn RN, Rabiner EA, Wells L, Dakin SG, et al. Translocator Protein as an Imaging Marker of Macrophage and Stromal Activation in Rheumatoid Arthritis Pannus. Journal of Nuclear Medicine. 2018;59:1125–1132. doi: 10.2967/jnumed.117.202200

101. Zhang HW, Xu AD, Sun X, Yang YQ, Zhang L, Bai H, Ben JJ, Zhu XD, Li XY, Yang Q, et al. Self-Maintenance of Cardiac Resident Reparative Macrophages Attenuates Doxorubicin-Induced Cardiomyopathy Through the SR-A1-c-Myc Axis. Circ Res. 2020;127:610–627. doi: 10.1161/Circresaha.119.316428

102. Kashiyama N, Miyagawa S, Fukushima S, Kawamura T, Kawamura A, Yoshida S, Harada A, Watabe T, Kanai Y, Toda K, et al. Development of PET Imaging to Visualize Activated Macrophages Accumulated in the Transplanted iPSc-Derived Cardiac Myocytes of Allogeneic Origin for Detecting the Immune Rejection of Allogeneic Cell Transplants in Mice. Plos One. 2016;11. doi: ARTN e0165748. 10.1371/journal.pone.0165748

103. Thai PN, Daugherty DJ, Frederich BJ, Lu XY, Deng WB, Bers DM, Dedkova EN, Schaefer S. Cardiac-specific Conditional Knockout of the 18-kDa Mitochondrial Translocator Protein Protects from Pressure Overload Induced Heart Failure. Sci Rep-Uk. 2018;8. doi: ARTN 16213. 10.1038/s41598-018-34451-2

104. Batarseh A, Papadopoulos V. Regulation of translocator protein 18 kDa (TSPO) expression in health and disease states. Mol Cell Endocrinol. 2010;327:1–12. doi: 10.1016/j.mce.2010.06.013

105. Valdes Olmos RA, Ten Bokkel Huinink WW, Dewit LG, Hoefnagel CA, Liem IH, vanTinteren H. Iodine-123 metaiodobenzylguanidine in the assessment of late cardiac effects from cancer therapy. Eur J Nucl Med. 1996;23:453–458. doi: Doi 10.1007/Bf01247376

106. dos Santos MJ, da Rocha ET, Verberne HJ, da Silva ET, Aragon DC, Soares J. Assessment of late anthracycline-induced cardiotoxicity by I-123-mIBG cardiac scintigraphy in patients treated during childhood and adolescence. J Nucl Cardiol. 2017;24:256–264. doi: 10.1007/s12350-015-0309-y

107. Heidenreich PA, Bozkurt B, Aguilar D, Allen LA, Byun JJ, Colvin MM, Deswal A, Drazner MH, Dunlay SM, Evers LR, et al. 2022 AHA/ACC/HFSA Guideline for the Management of Heart Failure A Report of the American College of Cardiology/American Heart Association Joint Committee on Clinical Practice Guidelines. Journal of the American College of Cardiology. 2022;79:E253–E421. doi: 10.1016/j.jacc.2021.12.012

108. Cardinale D, Colombo A, Lamantia G, Colombo N, Civelli M, De Giacomi G, Rubino M, Veglia F, Fiorentini C, Cipolla CM. Anthracycline-Induced Cardiomyopathy Clinical Relevance and Response to Pharmacologic Therapy. Journal of the American College of Cardiology. 2010;55:213–220. doi: 10.1016/j.jacc.2009.03.095

109. Hoffmann DB, Fraccarollo D, Galuppo P, Frantz S, Bauersachs J, Tillmanns J. Genetic ablation of fibroblast activation protein alpha attenuates left ventricular dilation after myocardial infarction. Plos One. 2021;16. doi: ARTN e0248196. 10.1371/journal.pone.0248196

110. Sun YX, Ma MQ, Cao DD, Zheng AC, Zhang YY, Su Y, Wang JF, Xu YH, Zhou M, Tang YS, et al. Inhibition of Fap Promotes Cardiac Repair by Stabilizing BNP. Circ Res. 2023;132:586–600. doi: 10.1161/Circresaha.122.320781

111. Rosenkrans ZT, Massey CF, Bernau K, Ferreira CA, Jeffery JJ, Schulte JJ, Moore M, Valla F, Batterton JM, Drake CR, et al. [Ga-68]Ga-FAPI-46 PET for non-invasive detection of pulmonary fibrosis disease activity. Eur J Nucl Med Mol I. 2022;49:3705–3716. doi: 10.1007/s00259-022-05814-9

112. Shi JL, Hou ZL, Yan J, Qiu WF, Liang LX, Meng MY, Li L, Wang XD, Xie YH, Jiang LH, Wang WJ. The prognostic significance of fibroblast activation protein-u in human lung adenocarcinoma. Ann Transl Med. 2020;8. doi: ARTN 224. 10.21037/atm.2020.01.82

